# Stereociliary localization and function of the tip-link protein PCDH15-CD2 are regulated by the PDZ-scaffold NHERF-1

**DOI:** 10.64898/2026.10.27.762349

**Authors:** Aray Adylkhan, Anna Andreeva, Xufeng Qiu, Zariat Shatkenova, Phillip Wilson, Alison Lim, Daniel Son, Raymond Huang, Ulrich Mueller, Xiaowei Lu

## Abstract

Sensory hair cells in the inner ear convert mechanical sound deflections into electrical signals via the stereociliary hair bundle on their apical surface. Proper bundle function requires that structural, adhesion, and motor proteins be delivered to specific stereocilia sub-compartments and maintained at precise levels throughout life, yet the underlying mechanisms remain poorly understood. Here we investigated how spatiotemporal localization of Protocadherin-15 (PCDH15), a component of kinociliary links, transient lateral links, and tip links, is regulated in mouse auditory hair cells. Using biochemical assays *in vitro*, we identified the PDZ-domain scaffolds NHERF-1/2 as direct binding partners of CD2, a PCDH15 isoform essential for hearing. We mapped the interaction to NHERF-1’s PDZ1 domain and CD2’s PDZ-binding motif (PBM), and showed that this binding is regulated by phosphorylation of the CD2 PBM. Using *Nherf1* knockout and *Pcdh15-CD2* knock-in mice defective in this interaction, we found CD2-NHERF-1 binding is required for proper CD2 levels and localization in the developing hair bundle. Loss of this interaction largely spared early bundle morphogenesis and MET function but caused high-frequency hearing loss and degeneration of the shortest stereociliary row in adult mice. Together, these findings define an important regulatory mechanism with therapeutic relevance for hair bundle repair.

## Introduction

Our sense of hearing relies on mechanosensory hair cells in the cochlea, which convert mechanical sound vibrations into electrical signals through a process called mechanoelectrical transduction (MET). Essential for their MET function, hair cells project actin-based protrusions that form the stereociliary hair bundle on their apical surface^1^. The hair bundle consists of actin-filled stereocilia organized in a V-shaped staircase pattern, with its vertex pointing to the lateral side of the cochlear duct. These are hallmark features of planar cell polarity, or polarity along the tissue plane, which is essential for the directional sensitivity of the hair bundle to sound stimuli.

Hair bundle formation begins with the centrifugal migration of the kinocilium, a specialized primary cilium, from the center to the lateral pole of the hair cell apex. This is followed by graded stereocilia elongation and formation of multiple types of extracellular links essential for hair bundle structural cohesion and function^2,3^. In particular, the cadherin superfamily members Protocadherin-15 (PCDH15) and Cadherin-23 (CDH23) are components of several types of extracellular links, including transient lateral links and tip links that interconnect neighboring stereocilia, and kinociliary links that tether the tallest stereocilia to the kinocilium at the vertex of the V-shaped hair bundle^4–10^. These link complexes have distinct functions during hair bundle development and maturation. Present only during development, transient lateral links maintain hair bundle cohesion, while kinociliary links establish proper orientation and the characteristic V-shape of the hair bundle^3,7^. Mutations in *Pcdh15* and *Cdh23* cause Usher syndrome type 1, the most common inherited cause of human deaf-blindness^11,12^. In mature hair cells, tip links extend from the tips of shorter stereocilia, where MET channels are localized, to adjacent taller stereocilia, mechanically gating these channels to enable hearing^3–5,7,9,10,13^. Despite their functional importance, little is known about the mechanisms that precisely regulate the amount and localization of PCDH15 and CDH23 proteins in stereocilia during hair bundle development and maintenance.

During development, alternative splicing generates three major PCDH15 cytoplasmic domain (CD) isoforms, CD1, CD2, and CD3, which share identical extracellular and transmembrane domains but contain distinct cytoplasmic tails^8^. Knockout of single *CD* isoforms in mice showed CD1 and CD3 are individually dispensable for hair bundle development and function, whereas loss of CD2 alone causes deafness, loss of kinociliary links, and disrupted hair-bundle planar polarity^10^. Conditional postnatal deletion of CD2 showed CD2 is specifically required to maintain tip links in mature hair cells, even though CD1/CD3 can substitute during development^4^. Thus, proper spatiotemporal localization of CD2 to distinct hair bundle compartments is essential for its multiple functions: forming transient lateral links along the stereocilia shafts, extending from the kinocilium to form kinociliary links, and partnering with the MET complex at the tips of shorter stereocilia in mature hair cells^14,15^. While it is shown that kinociliary CD2 targeting is mediated through a FGFR1-dependent intraflagellar transport pathway^13^, the mechanisms that regulate the level and spatial distribution of CD2 within stereocilia remain poorly understood.

To identify regulators of stereociliary localization of CD2, we screened for CD2 binding partners using affinity purification followed by mass spectrometry. This approach identified PDZ-domain scaffold protein Na^+^/H^+^ exchanger regulatory factors 1 and 2 (NHERF-1 and NHERF-2) as novel CD2 interactors. We demonstrated that the first PDZ (PDZ1) domain of NHERF-1 directly binds the C-terminal PDZ-binding motif (PBM) of CD2, and their binding affinity is regulated by phosphorylation of multiple serine and threonine residues in and around the CD2 PBM. We determined the *in vivo* role of the CD2-NHERF-1 interaction using two complementary mouse models: *Nherf1* knockout mice and *Pcdh15-CD2* knock-in mice with mutated PBM phosphorylation sites. We found that disrupted CD2-NHERF-1 interaction reduced CD2 levels and altered its distribution in the developing hair bundle. Using scanning electron microscopy (SEM) and patch-clamp recordings, we showed that hair bundle planar polarity and MET function are largely preserved in *CD2* knock-in mice deficient for NHERF-1 binding. However, adult *CD2* mutant mice develop high-frequency hearing loss accompanied by degeneration of the shortest (third-row) stereocilia. Together, our findings identify NHERF-1 as a scaffold for CD2 stereociliary localization and reveal a PDZ–PBM-dependent mechanism important for PCDH15-CD2 function in hearing.

## Results

### NHERF-1 is a Novel Direct Binding Partner of PCDH15-CD2

To identify novel binding partners of PCDH15-CD2 (CD2 hereafter), we performed an *in vitro* affinity pull-down assay coupled with tandem mass spectrometry (MS) analysis. Specifically, we used the cytoplasmic domain of CD2 (CD2-Cyto) fused to maltose-binding protein (MBP) as bait to capture interacting proteins from P0 mouse brain and lung tissue lysates. MBP alone served as a negative control for non-specific binding proteins. This approach identified two related PDZ domain-containing scaffolding proteins, NHERF-1 and NHERF-2, as novel CD2 binding partners (Fig.1A). NHERF proteins are well-established PDZ scaffold proteins linking membrane proteins with cytoskeletal proteins like Ezrin, Radixin, and Moesin (ERM) proteins, which were also identified in our pull-down experiment (Fig. 1A).

**Figure 1.**
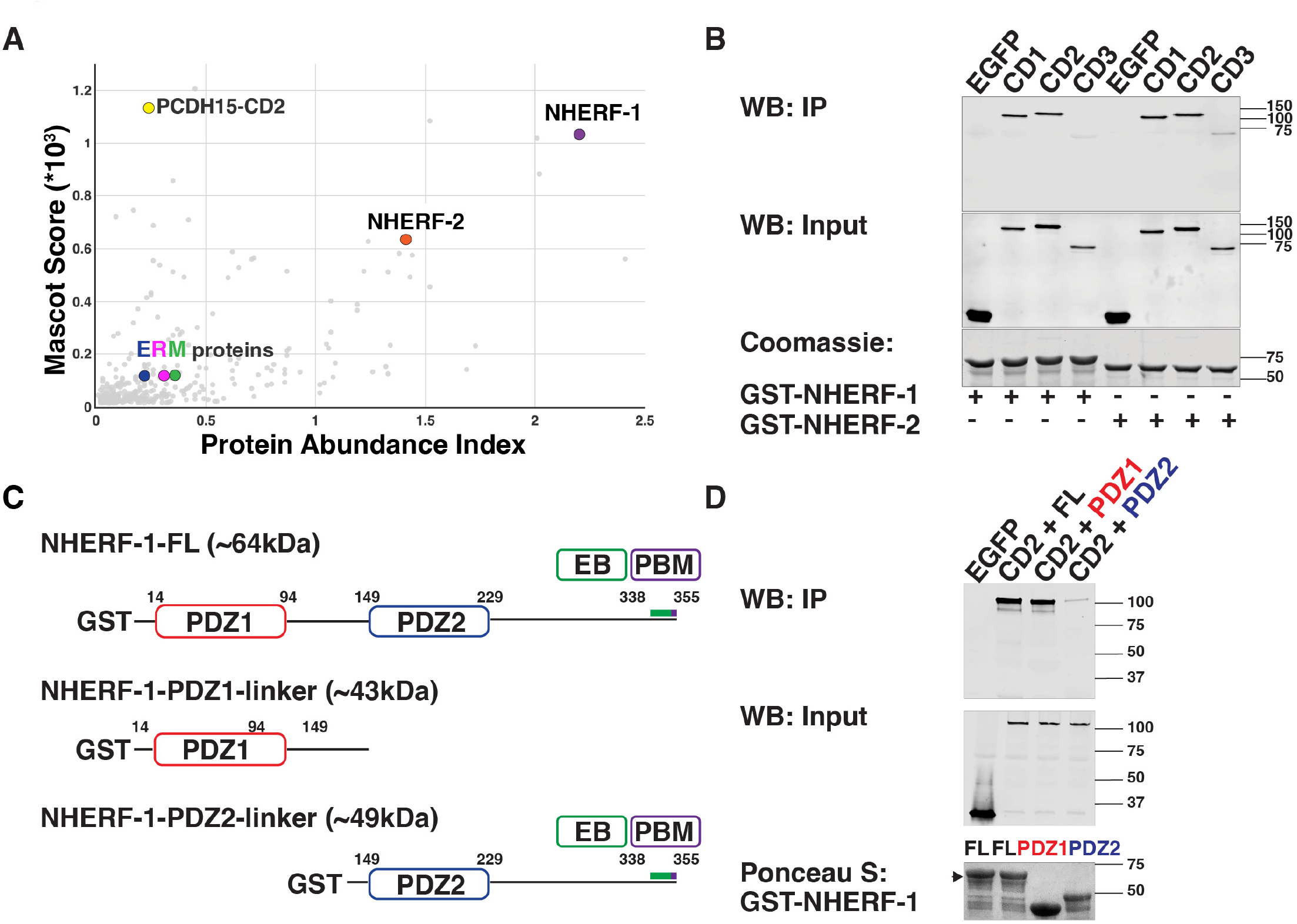
NHERF-1 and NHERF-2 are novel binding partners of PCDH15-CD2. **(A)** PCDH15-CD2 interactome identified by affinity purification - mass spectrometry (see Methods for calculations of Mascot Score and Protein Abundance Index). Dots represent specific proteins associated with His-MBP-CD2 Cyto, but not His-MBP; NHERF-1 and NHERF-2 are highly significant CD2 interactors. **(B)** GST-NHERF-1/2 pull-down of PCDH15 CD isoforms. Western blot analysis of bound fractions (top panel), whole-cell extracts from HEK293 cells overexpressing EGFP, EGFP-CD1, -CD2, and -CD3 (TM+Cyto) (middle panel), and Coomassie staining for GST-NHERF-1/2 bait proteins (bottom panel). **(C)** Schematic diagrams of NHERF-1 domains and GST-NHERF-1 truncation constructs used in D. EB, ERM-binding. **(D)** The PDZ1 but not PDZ2 domain of NHERF-1 mediates CD2 binding. Western blot analysis of bound fractions (top panel), whole-cell extracts from HEK293 cells overexpressing EGFP or EGFP-CD2 (TM+Cyto) (middle panel), and Ponceau staining for GST-NHERF-1 full-length (FL) and truncation proteins (bottom panel).

To test for direct binding between NHERF-1/NHERF-2 and the three major PCDH15 isoforms (CD1, CD2 and CD3), we first expressed each of the EGFP-tagged CD1, CD2 or CD3 containing the transmembrane and cytoplasmic domains (TM+Cyto)^13^ in HEK 293T cells, and incubated the respective cell lysates with immobilized GST-NHERF-1 and GST-NHERF-2 proteins *in vitro* (Fig. 1B). EGFP-expressing cell lysates served as negative control. Western blot analysis demonstrated that NHERF-1 and NHERF-2 bind with high affinity to CD1 and CD2, but not CD3. We next focused on NHERF-1 because previous studies have demonstrated its role in hair bundle morphogenesis and hearing; in contrast, *Nherf2* knockout mice exhibit normal hearing^16^. The C-terminus of CD2 contains a canonical Class I PDZ-binding motif (PBM) (-X-S/T-X-V/L), while NHERF-1 contains two tandem PDZ domains (PDZ1 and PDZ2, Fig. 1C), suggesting CD2-NHERF-1 binding is mediated through PDZ-PBM interactions. To test this, we performed GST pulldown assays using either PDZ1- or PDZ2-containing NHERF-1 fragments as bait, and found that the PDZ1 but not PDZ2 domain of NHERF-1 was both necessary and sufficient to mediate CD2 binding (Fig. 1D). Because NHERF proteins are highly homologous, NHERF-2 likely also binds CD2 PBM through its PDZ1 domain.

### Role of NHERF-1 in Planar Polarity and Hair Bundle Morphogenesis

Having identified NHERF-1/2 as novel CD2 binding partners, we next investigated the physiological relevance of this interaction *in vivo*, using previously characterized global *Nherf1* knockout (*Nherf1^KO/KO^*) mice^16,17^. First, we analyzed NHERF-1 expression in the organ of Corti (OC) during early postnatal development. In wild-type (*Nherf1^WT^*) hair cells at postnatal day 0 (P0), NHERF-1 is localized along the length of stereocilia, with higher levels in outer hair cells (OHCs) than inner hair cells (IHCs) (Fig. 2A, C; Fig. S1A). By P5, its localization in the OHC hair bundle becomes restricted to a middle zone of stereocilia, below the tips of the tallest row of stereocilia, and is barely detectable in IHCs (Fig. 2B, C; Fig. S1B). Interestingly, NHERF-1 levels in OHC hair bundles exhibit a base-to-apex increasing gradient, which is more pronounced at P5 than at P0 (Fig. S1A, B). In addition, NHERF-1 is highly expressed on the apical surface of supporting cells at both stages, where its level significantly increased by P5 (Fig. 2B, C; Fig. S1A, B). NHERF-1 staining was lost in the *Nherf1^KO/KO^* cochlea, confirming antibody specificity (Fig. 2A, B).

**Figure 2.**
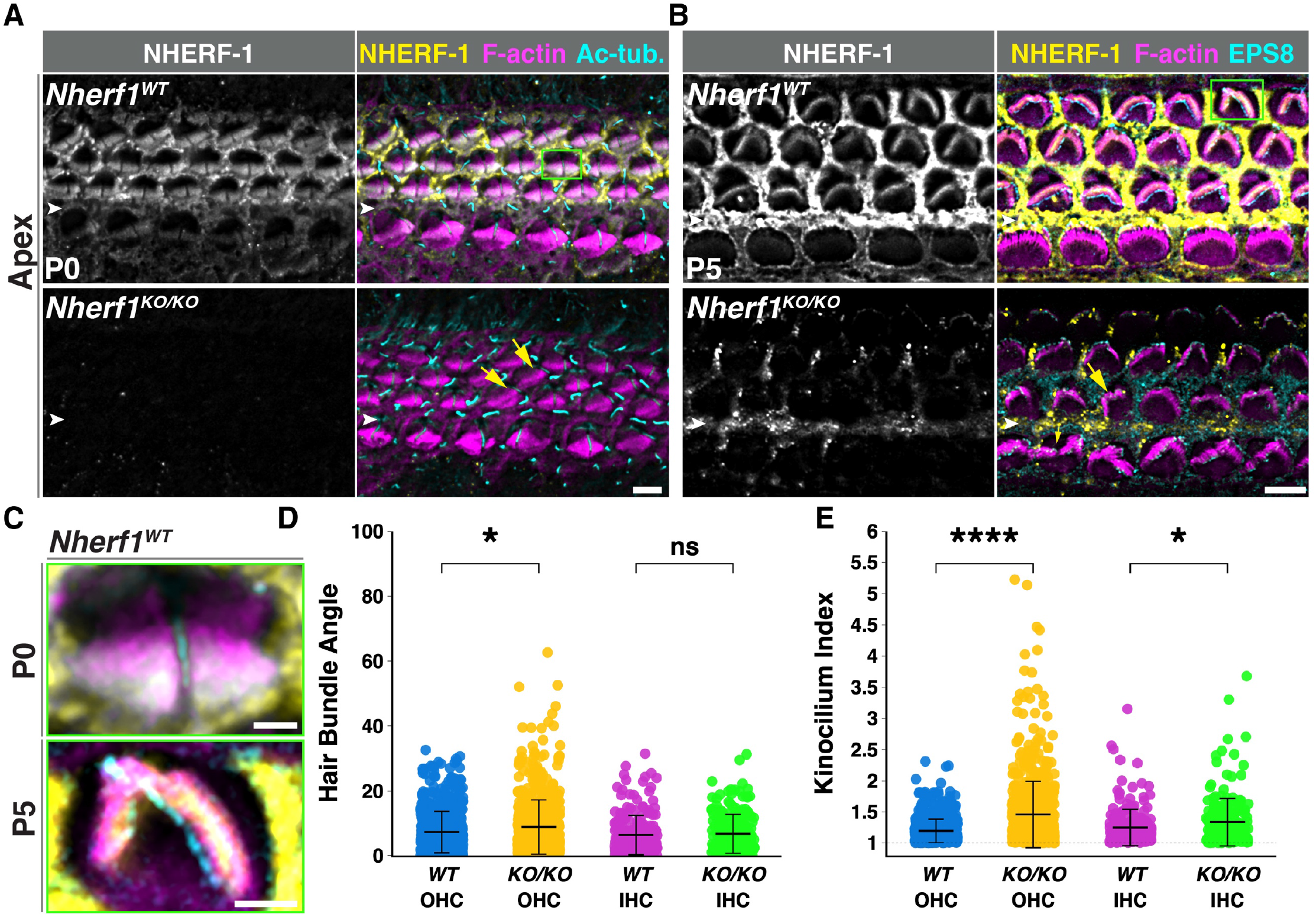
NHERF-1 localization in the organ of Corti and hair cell planar polarity defects in *Nherf1^KO/KO^* mice. **(A, B)** Whole-mount organ of Corti surface preparations stained for NHERF-1 (yellow), F-actin (magenta), acetylated tubulin (Ac-tub, cyan in A), or EPS8 (cyan in B) at P0 (A) and P6 (B). Yellow arrows indicate misoriented hair bundles in *Nherf1^KO/KO^*. **(C)** Magnified views of selected *Nherf1^WT^* OHCs in A and B (indicated by green boxes). White arrowheads mark the inner pillar cell (IPC) row. Scale bars: A, B, 6 μm; C, 2 μm. **(D, E)** Quantification of hair bundle orientation (D) and kinocilium index (E) at P0. The statistical test uses a linear mixed-effects model (N = 4 independent experiments). Data presented as mean ± SD. *,p≤0.02; ****P ≤ 0.0001; ns, non-significant.

To determine NHERF-1’s role in hair cell planar polarity, which is regulated by CD2, we analyzed hair bundle orientation and kinocilium positioning in *Nherf1^KO/KO^* hair cells at P0. As hallmarks of hair cell planar polarity, WT hair cells each have a chevron-shaped hair bundle with their vertices all pointing laterally; the kinocilium is tethered to the vertex of the hair bundle. In *Nherf1^KO/KO^* OC, OHCs but not IHCs displayed significant hair bundle misorientation (Fig. 2A, yellow arrows; Fig. 2D). Although we did not observe detachment of the kinocilium from the hair bundle, many hair bundles of OHCs, and to a lesser extent, IHCs, had an off-center kinocilium (kinocilium index > 1.5, see methods) (Fig. 2A, 2E). These hair cell planar polarity defects persisted at P5, and many IHC and OHC hair bundles in *Nherf1^KO/KO^* were disorganized and/or misshapen, albeit still with a recognizable staircase pattern. Thus, these results indicate a role of NHERF-1 in regulating hair cell planar polarity and hair bundle morphogenesis.

### NHERF-1 is Required for the Localization of PCDH15-CD2 in Stereocilia

To investigate the specific role of NHERF-1 in CD2 regulation during hair bundle morphogenesis, we asked whether NHERF-1 is required for CD2 localization *in vivo*. Consistent with previous studies^10^, CD2 is highly enriched in the P0 OHC hair bundle. In stark contrast, CD2 levels were significantly reduced in the *Nherf1^KO/KO^* OHC hair bundle at all cochlear positions; the residual CD2 was localized towards the stereocilia tips (Fig. 3, Fig. S2). To assess the overall level and distribution of PCDH15, we stained P0 cochleae with pan-PCDH15 antibodies, which detect all PCDH15 isoforms^18^. We found that total levels of PCDH15 in *Nherf1^KO/KO^* OHC hair bundles were also reduced compared to the *Nherf1^WT^*, albeit to a lesser degree than CD2 (Fig. S3). These results suggest that stereociliary localization of CD3^10^, which exhibits low binding affinity for NHERF-1 *in vitro*, was not impacted in *Nherf1^KO/KO^* hair cells. Because Cadherin-23, which partners with PCDH15 to form extracellular links, was shown to bind NHERF-1 *in vitro*^5,16^, we also examined the localization of Cadherin-23^19^. In contrast to CD2, Cadherin-23 is localized to both *Nherf1^WT^* and *Nherf1^KO/KO^* hair bundles at comparable levels at P0 (Fig. S4). Taken together, these results demonstrate that NHERF-1 is specifically required for the proper level and localization of CD2 in the developing hair bundle *in vivo*.

**Figure 3.**
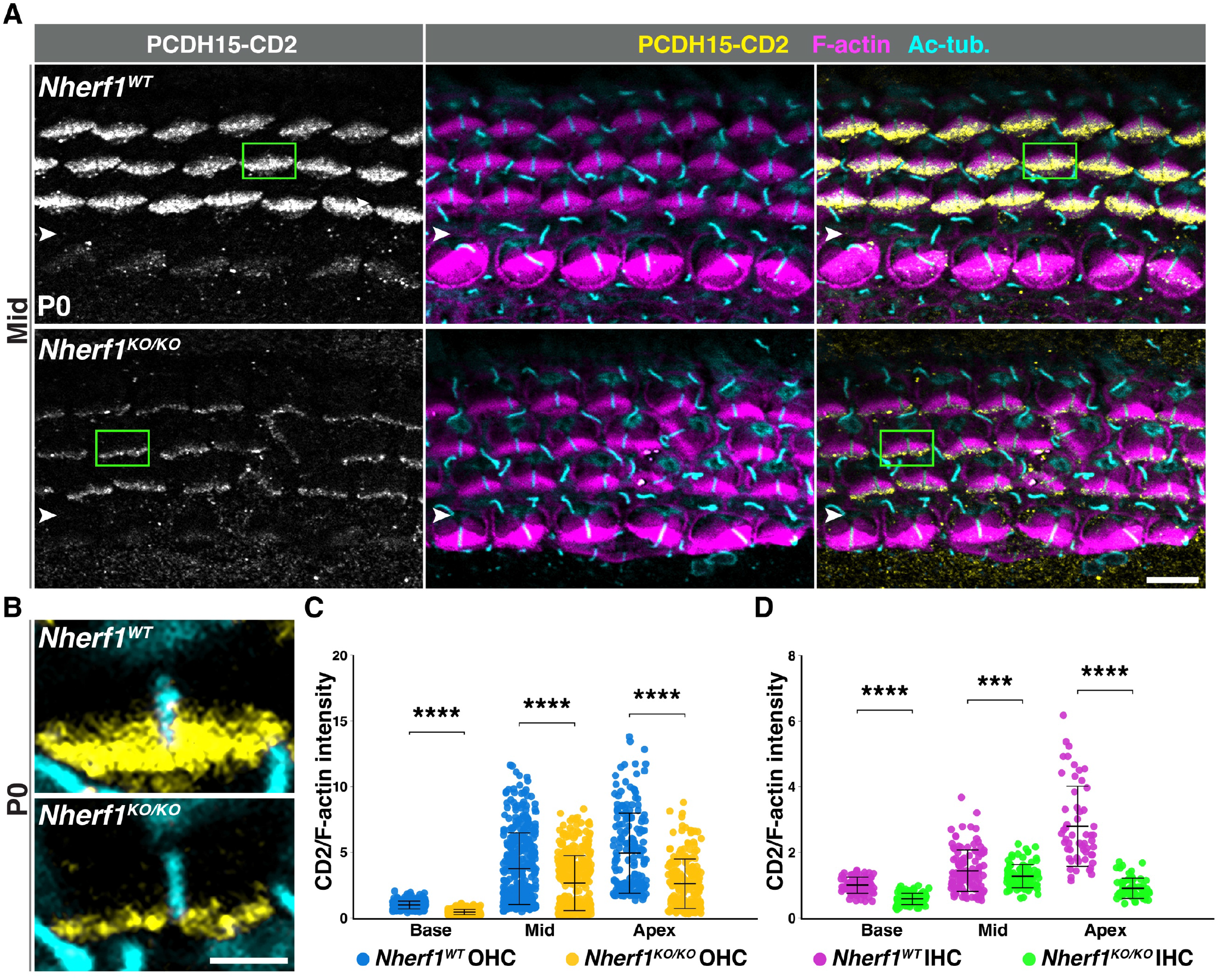
Reduced PCDH15-CD2 staining in OHC hair bundles of *Nherf1^KO/KO^* mice at P0. **(A)** PCDH15-CD2 staining (yellow) in *Nherf1^WT^* and *Nherf1^KO/KO^* cochleae, counterstained for F-actin (magenta) and acetylated tubulin (Ac-tub, cyan) at P0. **(B)** Magnified views of selected OHCs from A and B (indicated by green boxes. White arrowheads mark the IPC row. Scale bars: A, 6 μm; B, 2 μm. (**C, D**) Quantification of normalized PCDH15-CD2 staining intensity in OHCs (C) and IHCs (D) at indicated cochlear positions. The statistical test uses a linear mixed-effects model (N = 3 independent experiments). Data are presented as mean ± SD. ***P ≤ 0.001; ****P ≤ 0.0001.

### The CD2-NHERF-1 Interaction is Modulated by CD2 Phosphorylation

It is well established that PDZ-PBM interactions can be regulated by phosphorylation of residues within or adjacent to the PBM^20^. To further investigate how CD2 stereociliary localization is regulated, we explored the possibility that the CD2-NHERF-1 interaction is modulated by phosphorylation of the CD2 cytoplasmic tail. To this end, we first asked whether EGFP-tagged CD2 (TM+Cyto) overexpressed in HEK293T cells is phosphorylated. Using affinity pull-down followed by MS analysis, we identified three phosphorylation sites in the CD2 cytoplasmic tail region: S1771, S1776, and T1781 (Fig. 4A-C). To gain structural insight into the role of these residues in the CD2-NHERF-1 binding interaction, we performed AlphaFold modeling, which predicted a high-confidence complex between the NHERF-1 PDZ1 domain and the C-terminal PBM of CD2, in agreement with our experimental data (Fig. S5A, B). In the AlphaFold model, the hydroxyl group of T1781 in the CD2 PBM forms a critical hydrogen bond with H59 in the canonical binding groove of the PDZ1 of NHERF-1 (Fig. S5B), while S1771 and S1776 lie immediately N-terminal to the PBM and could make surface contacts with the PDZ1. Interestingly, the AlphaFold model predicts that both phosphorylation of T1781 and a phospho-deficient mutation T1781A disrupt the hydrogen bond formation: the former via steric hindrance and the latter through the loss of the hydroxyl group. (Fig. S5C, D).

**Figure 4.**
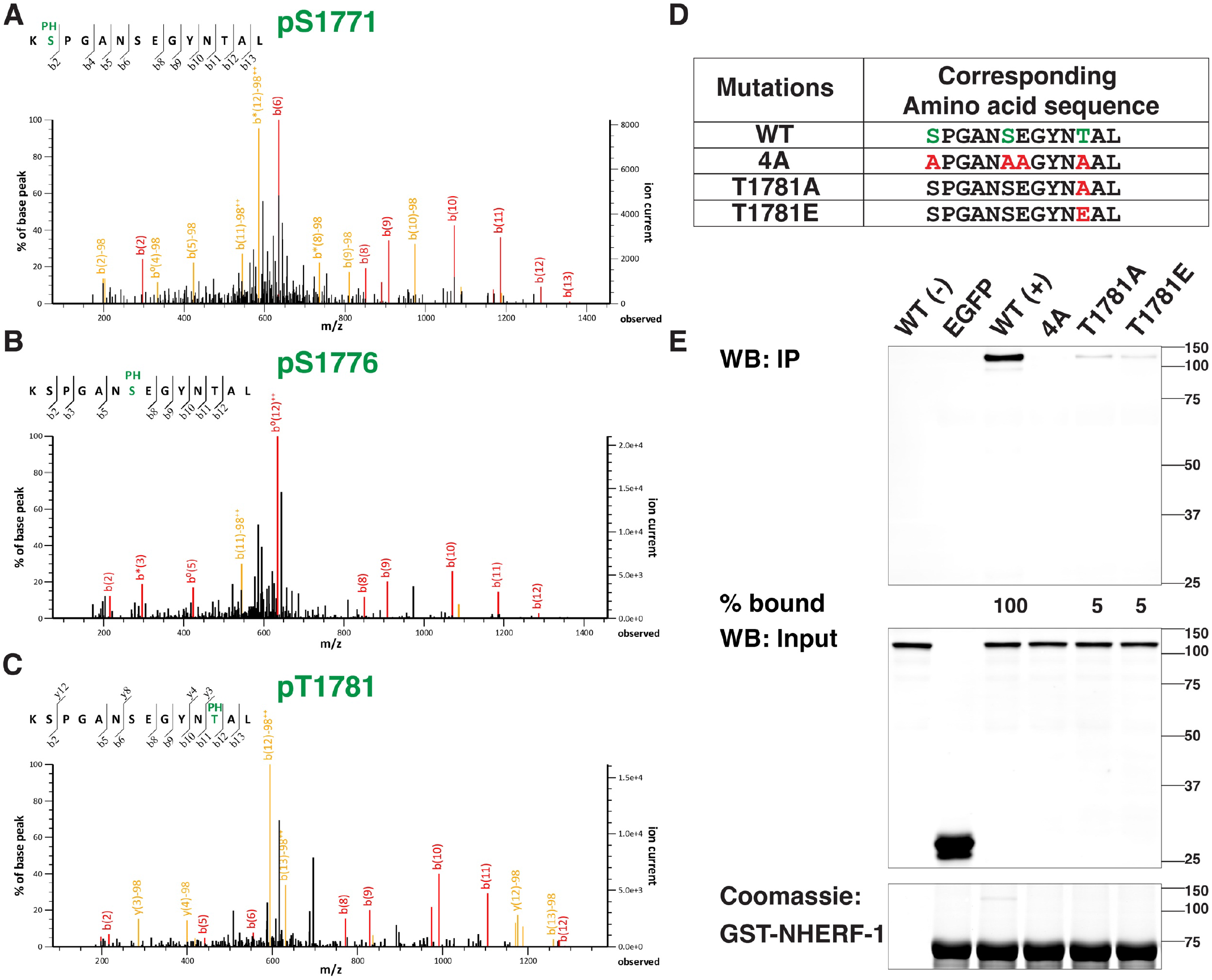
CD2-NHERF-1 binding is modulated by phosphorylation of the CD2 cytoplasmic tail. **(A–C)** Analysis of MS/MS fragmentation spectra of the PCDH15-CD2 cytoplasmic tail identified phosphorylation at S1771 (A), S1776 (B), and T1781(C). Relevant b and y ions are labeled according to the accepted nomenclature ^51^. **(D)** C-terminus sequences of wild-type (WT) and various mutant EGFP-CD2 (TM+Cyto) constructs. **(E)** Mutations of CD2 phosphorylation sites impair NHERF-1 binding *in vitro*. Western blot analysis of CD2 bound to GST-NHERF-1 (top panel), whole-cell extracts from HEK293 cells overexpressing EGFP, wild-type and indicated mutants of EGFP-CD2 (TM+Cyto) (middle panel), and Coomassie staining for GST-NHERF-1 bait proteins (bottom panel).

To determine whether phosphorylation of T1781, as well as S1771 and S1776, influences CD2-NHERF-1 binding affinity, we generated single and triple phospho-deficient alanine substitution mutant constructs, CD2-T1781A and CD2-4A, respectively. CD2-4A carries an additional E1777A mutation, to further neutralize the negative charge of the extended PBM region (Fig. 4D). In GST pulldown assays, CD2-NHERF-1 binding was greatly reduced by the T1781A and abolished by the CD2-4A mutations (Fig. 4E). To investigate the effect of T1781 phosphorylation on CD2-NHERF-1 interaction, we also generated a phospho-mimetic mutant construct, CD2-T1781E. In agreement with the AlphaFold prediction, the T1781E mutation significantly decreased CD2-NHERF-1 binding, to a similar extent as T1781A (Fig. 4E). Together, these results indicate that the CD2-NHERF-1 interaction is influenced by multiple phosphorylation sites in the CD2 cytoplasmic tail; T1781 phosphorylation negatively regulates CD2-NHERF-1 binding interactions, whereas phosphorylation at S1771 and S1776 likely acts as a positive regulator.

### *CD2^4A/4A^* Knock-in Mice Had Largely Normal Hair Bundle Morphogenesis, but Reduced CD2 Stereociliary Localization

Our earlier results indicate that NHERF-1 is required for CD2 stereociliary localization, likely by directly binding to CD2. The discovery that CD2-NHERF-1 binding is regulated by CD2 phosphorylation now provided us an opportunity to more specifically interrogate the physiological significance of this interaction *in vivo*. To this end, we generated a *CD2^4A/4A^* knock-in mouse model using CRISPR/Cas9. The *CD2^4A^* allele carries the equivalent four alanine substitution mutations that abolished NHERF-1 binding *in vitro* (Fig. 4D). We first asked whether any of the identified phosphorylation sites were phosphorylated *in vivo*. To this end, we attempted to generate phospho-specific antibodies against CD2 cytoplasmic tail individually phosphorylated at S1771, S1776, and T1781, respectively, and were only successful in obtaining phospho-S1771-specific antibodies (Fig. S6). Interestingly, CD2 phospho-S1771 signals were detected at the tips of both the kinocilium and stereocilia in *CD2^WT^* but not *CD2^4A/4A^* OHCs, indicating that CD2 is phosphorylated *in vivo* in the developing hair bundle (Fig. S6A).

Next, we analyzed hair cell planar polarity and hair bundle morphogenesis in *CD2^4A/4A^* cochleae at P0 and P6 (Fig. 5A, B). At P0, mild defects in hair bundle orientation and kinocilium positioning were observed, affecting OHCs and IHCs to different degrees (Fig. 5A, C, D). At P6, immunofluorescence and SEM analysis showed mild hair bundle orientation and shape defects, while hair bundle cohesion and staircase formation were largely intact (Fig. 5B, E). We then analyzed CD2 localization in *CD2^4A/4A^* hair cells. If NHERF-1 regulates CD2 stereociliary localization via a direct interaction, then CD2 localization in *CD2^4A/4A^* should phenocopy that of *Nherf1^KO/KO^*. This was indeed the case: along the entire cochlear length at P0, CD2 staining intensity was significantly reduced in *CD2^4A/4A^* OHC hair bundles and only detectable towards the stereocilia tips (Fig. 6A, C and Fig. S7). Moreover, pan-PCDH15 staining showed a reduction of total PCDH15 levels in stereocilia comparable to *Nherf1^KO/KO^* (Fig. S8), suggesting that CD1/CD3 isoforms were not significantly upregulated in *CD2^4A/4A^* hair cells at P0. We also analyzed the localization of Cadherin-23 and NHERF-1, and found that their levels and localization in *CD2^4A/4A^* hair bundles were comparable to *CD2^WT^* (Fig. S9). As CD2 is the only isoform transported into the kinocilium for kinociliary links^10^, we used super-resolution microscopy to resolve CD2 localization in kinocilium, and found that CD2 staining along the kinocilium appeared intact in *CD2^4A/4A^* OHCs, suggesting that the CD2-4A mutation does not affect kinociliary links (Fig. 6B). Together, these data indicate that disruption of CD2-NHERF-1 interaction *in vivo* significantly decreases stereociliary localization of CD2 but not Cadherin-23 or NHERF-1, resulting in mild hair cell planar polarity and hair bundle shape defects.

**Figure 5.**
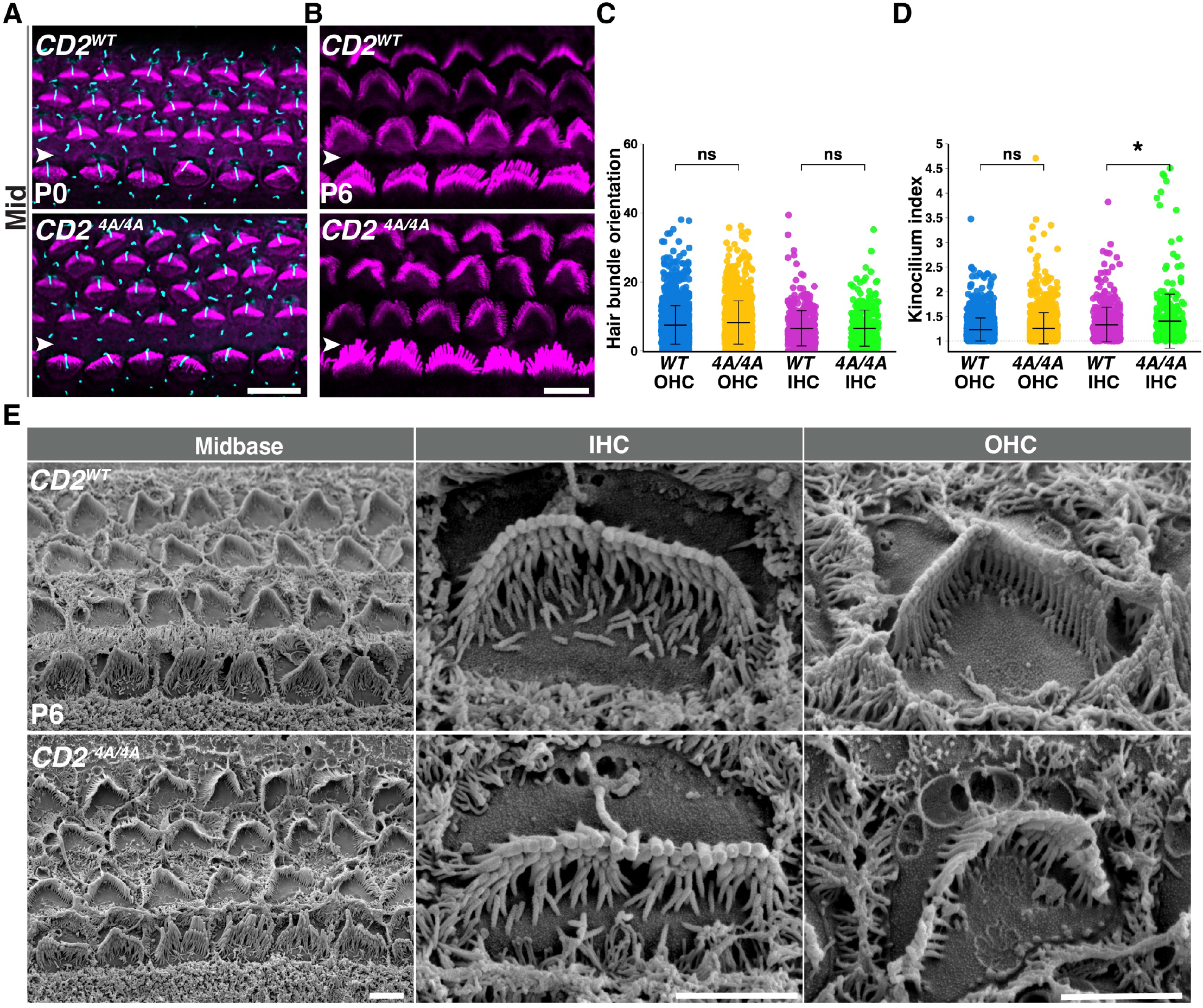
Mild defects in planar polarity and hair bundle morphology in *CD2 ^4A/4A^* hair cells. **(A, B)** Whole-mount surface preparations of the middle turn of the organ of Corti stained for F-actin (magenta) and acetylated tubulin (cyan) at P0 (A) and P5 (B). White arrowheads mark the IPC row. **(C, D)** Quantification of hair bundle orientation (C) and kinocilium index (D) at P0. Statistical test using a linear mixed-effects model (N=3 independent experiments). Data presented as mean ± SD. *, p≤0.02; ns, non-significant. **(E)** Representative SEM images of the midbasal region of the organ of Corti and individual IHCs and OHCs at P6. Scale bars, A-B, 6 μm; E, 2 μm.

**Figure 6.**
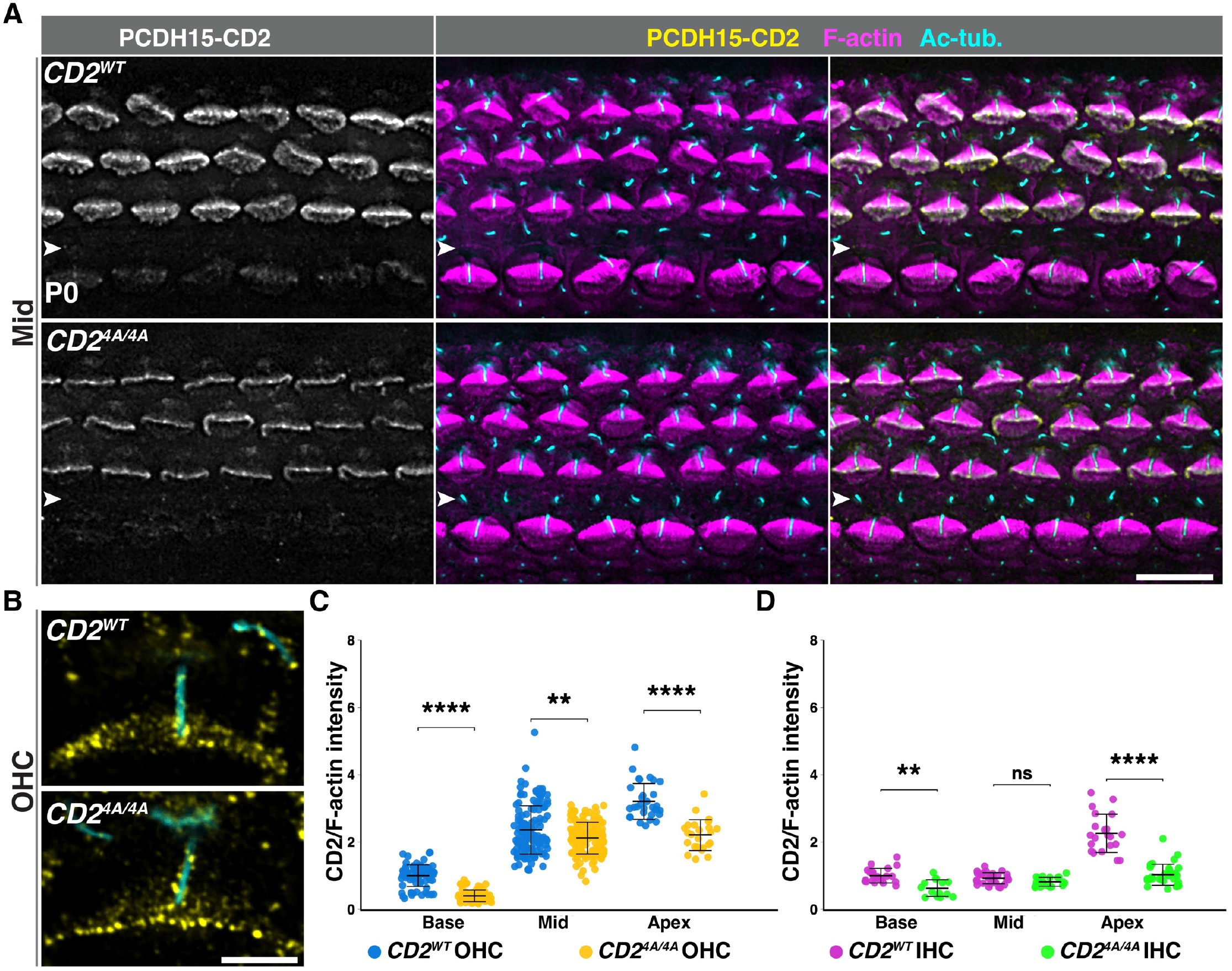
Reduced PCDH15-CD2 localization in OHC hair bundles of *CD2 ^4A/4A^* mice at P0. **(A)** Whole-mount organ of Corti surface preparations from the middle turn stained for PCDH15-CD2 (yellow), F-actin (magenta), and acetylated tubulin (Ac-tub, cyan). White arrowheads mark the IPC row. **(B)** Subcellular localization of CD2 (yellow) in OHCs at P0 revealed by super-resolution microscopy. The kinocilium is labeled by acetylated tubulin staining (cyan). Scale bars, A, 6 μm; B, 2 μm. **(C, D)** Quantification of normalized PCDH15-CD2 staining intensity in OHCs (C) and IHCs (D) at indicated positions. The statistical test uses a two-way ANOVA. Data presented as mean ± SD; **P ≤0.01; ****P ≤ 0.0001; ns, non-significant.

### *CD2^4A/4A^* Early Postnatal Hair Cells Had Normal Mechanoelectrical Transduction

CD2 is crucial for the formation of the tip link^4^. At P6, when hair cells become competent for MET, we observed punctate CD2 staining in both OHC and IHC hair bundles, with reduced staining intensity in *CD2^4A/4A^* compared to the wild-type (Fig. 7A). To evaluate MET function of *CD2^4A/4A^* mice, we first performed FM1-43 dye uptake assays, which require functional MET channels^21^. Both control and *CD2^4A/4A^* OHCs internalized FM1-43 at P6, suggesting CD2-4A is capable of sustaining MET (Fig. 7B).

**Figure 7.**
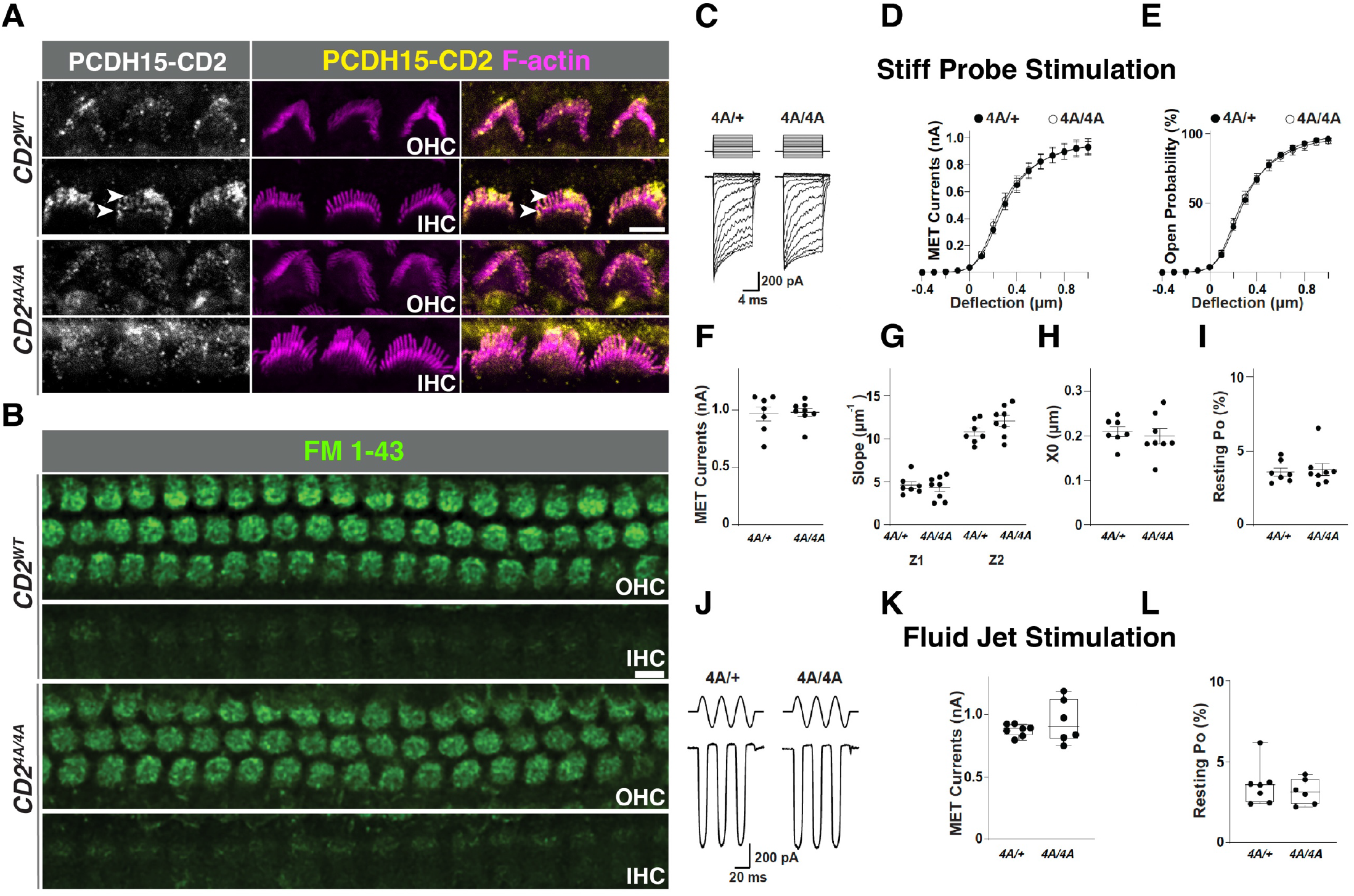
Normal MET recordings in early postnatal *CD2 ^4A/4A^* OHCs. **(A)** PCDH15-CD2 localization (yellow) in *CD2 ^WT^*and *CD ^4A/4A^* OHCs and IHC hair bundles (magenta) at P6. White arrowheads indicate the staining at the tallest and second row of stereocilia. **(B)** FM1-43 uptake in cochlear explants from the middle turn of *CD2 ^WT^* and *CD ^4A/4A^* at P6. Scale bars, A, B: 6 μm. **(C-E)** Representative MET currents (C), Current/displacement (I/X) (D), and open probability/displacement (Po/X) plots (E) of OHCs from the apical turn of *CD2^4A/+^* and *CD^4A/4A^* cochleae at P5-P6 stimulated with a stiff probe. **(F)** Peak MET currents derived from double Boltzmann fitting of I/X plots in D. **(G-I)** Slope (Z1 and Z2) plots (G), MET channel set point (X0) (H), and resting open probability (Po) (I) derived from the Po/X plots in E (n = 7 cells from *CD2^4A/+^* and 8 cells from *CD2^4A/4A^* mice). **(J-L)** Representative MET currents (J), peak MET current amplitude (K), and MET channel resting open probability (L) in response to sinusoidal fluid jet deflection of OHC hair bundles. (n = 7 OHCs from *CD2^4A/+^*, and 8 OHCs from *CD2^4A/4A^* mice). Two-sided unpaired t-tests showed no significant differences between the two genotypes.

To further assess the effect of CD2-4A mutations on tip link functions, we measured MET currents of OHCs in the mid-apical turn of the cochlea from *CD2^4A/+^* and *CD2^4A/4A^* mice at P5-P6. When stimulated by stepwise deflections with a stiff glass probe, the peak amplitudes of MET currents were not significantly different between *CD2^4A/+^* and *CD2^4A/4A^* OHCs (Fig. 7C, D, and F), suggesting that the total number of MET channels was not altered in *CD2^4A/4A^* mutants. Moreover, MET current/displacement (I/X) plots and MET channel open probability/displacement (Po/X) plots from *CD2^4A/+^* and *CD2^4A/4A^* OHCs (Fig. 7D, E) showed similar kinetics, with comparable slopes (Z1, Z2) and set point (X0) (Fig. 7G-I). Similarly, the resting open probability (P_o_) was also normal (Fig. 7I). We also observed normal MET currents and resting P_o_ when evoked by sinusoidal fluid-jet deflection of apical OHC hair bundles at P6-P7 (Fig. 7J-L). Together, these electrophysiological data indicate the CD2-4A mutations do not affect the force sensitivity or the gating kinetics of the MET channel in early postnatal hair cells. These findings are consistent with previous observations that early postnatal OHCs from global *Pcdh15-CD2* knockout mice also retained normal MET function^10^.

### *CD2^4A/4A^* Adult Mice Develop High-Frequency Hearing Loss

To determine the effect of disrupted CD2-NHERF-1 interaction on hearing, we performed Auditory Brainstem Response (ABR) testing on *CD2^4A/4A^* adult mice at multiple stages. At one month of age, *CD2^4A/4A^* mice displayed significantly elevated ABR thresholds across all frequencies tested, particularly at high frequencies (22-32 kHz) (Fig. 8A). Similar high-frequency hearing loss was observed at 2.5 and 5 months old. To understand the underlying cause of the hearing loss, we examined *CD2^4A/4A^* hair cells at P30 using SEM. There was no significant hair cell loss at this stage, indicating that hearing loss is not a result of hair cell degeneration. However, close examination revealed hair bundle structure defects in both *CD2^4A/4A^* IHCs and OHCs at the cochlear base, where high-frequency sounds are detected. In IHCs, we observed degeneration of the third-row stereocilia (Figure 8B, magenta). OHCs also showed mild stereocilia degeneration in the third row (Fig. 8B, asterisks). Therefore, we conclude that while CD2-NHERF-1/2 interaction is dispensable for MET function in the early postnatal period, it is essential for maintaining hair bundle morphology and hearing function in adult hair cells.

**Figure 8.**
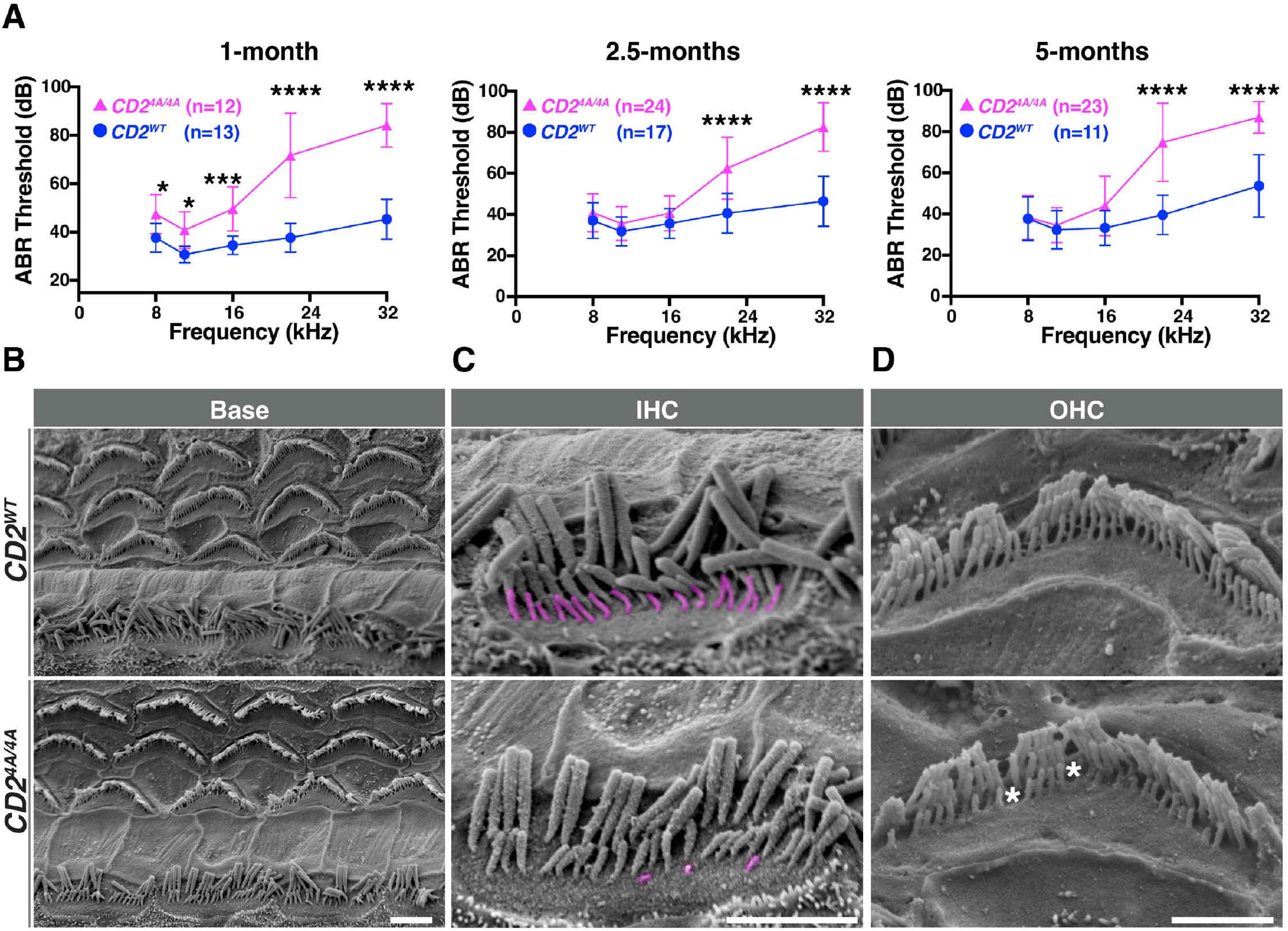
Adult *CD2^4A/4A^* mice develop high-frequency hearing loss accompanied by degeneration of the third-row stereocilia. **(A)** ABR thresholds in C*D2^WT^* (blue) and *CD2^4A/4A^*(magenta) mice at 1 month, 2.5 months, and 5 months, with the indicated number of animals. Data are presented as mean ± SD; Two-way ANOVA tests showed significant differences at 22kHz and 32KHz. * P ≤ 0.02; ***P ≤ 0.0002; ****P ≤ 0.0001. **(B-D)** SEM images of the organ of Corti from the basal turn of C*D2^WT^*and *CD2^4A/4A^* cochleae at P30 (B). High-magnification views of individual IHC (C) and OHC hair bundles (D) revealed degeneration of third-row stereocilia (pseudocolored in magenta) in *CD2^4A/4A^* IHCs and focal gaps in the third stereocilia row (asterisks) in OHCs. Scale bars, A, 1 μm; B, C, 0.5 μm.

## Discussion

The formation, maintenance, and function of the hair bundle require precise regulation of the amount and localization of hair bundle structural proteins within the stereocilia. However, little is known about mechanisms controlling proteostasis in the hair bundle compartment, particularly transmembrane hair bundle proteins. Our study identifies NHERF-1, a PDZ domain scaffold protein, as a novel regulator of PCDH15-CD2 level and localization in the developing hair bundle. We demonstrate that the PDZ1 domain of NHERF-1 directly binds the CD2 C-terminal PBM domain, and that this interaction is regulated by phosphorylation of the CD2 cytoplasmic tail and required for proper CD2 localization in the developing hair bundle. While *CD2^4A/4A^* knock-in mice deficient for NHERF-1 binding had largely normal hair bundle development and MET function at early postnatal stages, they developed high-frequency hearing loss and degeneration of the third row of stereocilia as young adults, indicating an essential role of this PDZ-PBM interaction in CD2 function in adult hair cells.

Our findings are consistent with NHERF-1 acting as a specific stereociliary targeting or anchoring scaffold for CD2. In *Nherf1^KO/KO^* hair cells, levels of CD2, but not CDH23, in the stereocilia were significantly decreased. Moreover, the distribution of CD2 within the stereocilia was shifted towards the tip. NHERF-1 may promote CD2 entry into stereocilia as a stereociliary targeting adaptor. Because there are no secretory vesicles within stereocilia, transmembrane hair bundle proteins are thought to be delivered to the plasma membrane in the apical periphery of hair cells, known as the pericuticular necklace, and then enter stereocilia by lateral diffusion^22^. MYO3A is capable of binding and transporting PCDH15-CD2 into filopodia *in vitro*^18,23^, however, whether myosin motors help transport PCDH15-CD2 into stereocilia *in vivo* is unknown. Since we did not observe CD2 accumulation in the pericuticular necklace region or intracellular vesicles of *Nherf1^KO/KO^* hair cells, as would be predicted by the targeting adaptor model, our results favor the alternative model wherein NHERF-1 acts as an anchoring scaffold for CD2 to capture and retain CD2 in stereocilia membranes (Fig. 9). This may be achieved by coupling CD2 to F-actin through ERM (Ezrin, Radixin, and Moesin) family proteins^24^. In support, ERM proteins were identified in our CD2 affinity pulldown experiments and have been shown to be required for normal hearing ^25,26^.

**Figure 9.**
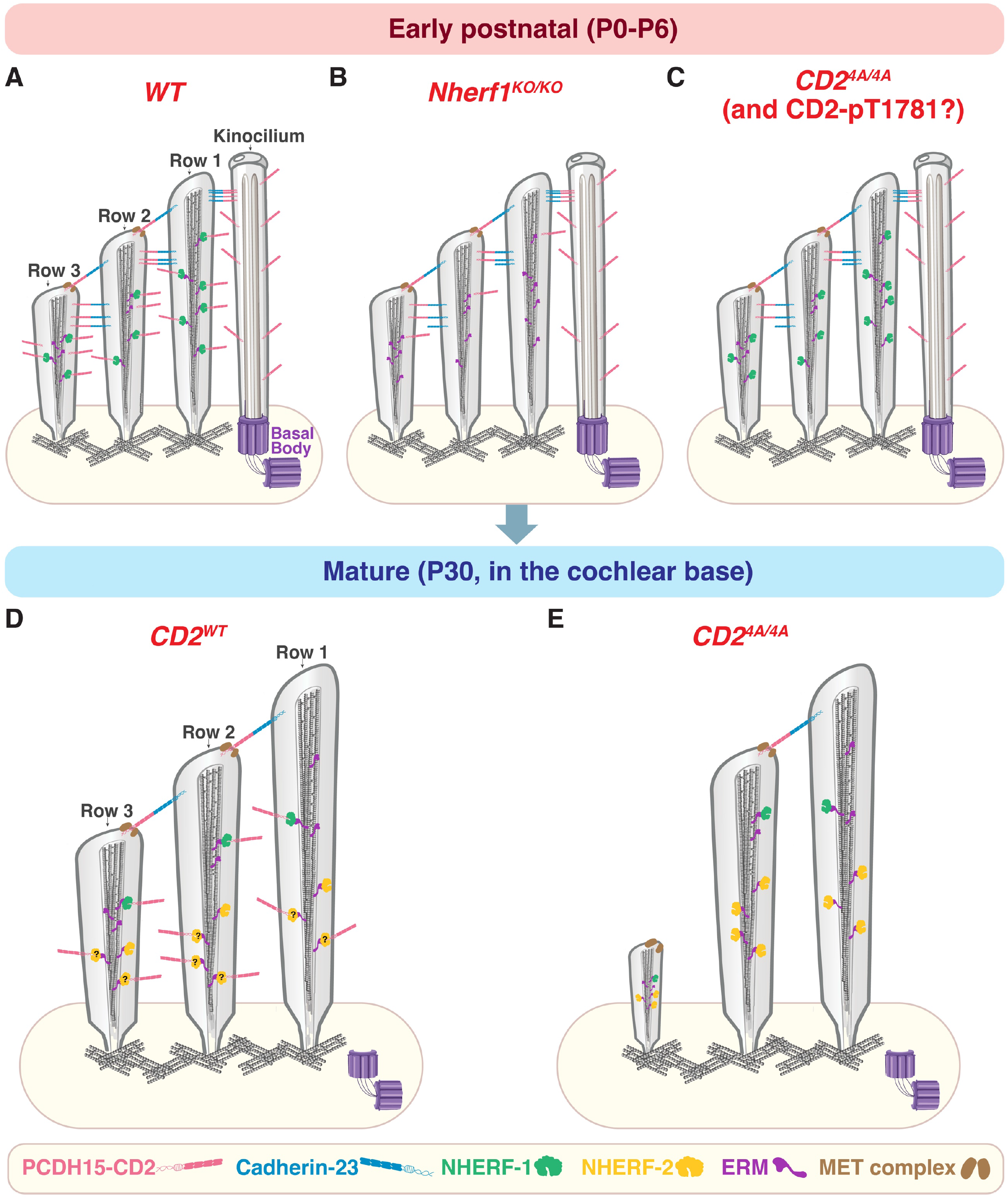
A Model for regulation of stereociliary localization of PCDH15-CD2 by NHERF-1/2 binding. **(A-C)** Early postnatal hair bundles (P0-P6). In *wild-type (WT)* hair bundles (A), NHERF-1 anchors CD2 in stereocilia membranes, likely through ERM-mediated coupling to F-actin. This interaction tunes the precise amount and localization of CD2 required to establish transient lateral links and later tip links. In *Nherf1^KO/KO^* (B) and in *CD2^4A/4A^* hair bundles (C), CD2 stereociliary localization is reduced. Nevertheless, remaining CD2 and other PCDH15 isoforms preserve early tip link formation and MET function in *CD2^4A/4A^*. Similarly, CD2 phosphorylated at T1781 is predicted to have reduced stereociliary localization. **(D, E)** Mature hair bundles (P30, in the cochlear base). In *WT* hair bundles (D), there is a pool of CD2 anchored in stereociliary membranes via NHERF-1/2, allowing for efficient tip-link maintenance and repair. In *CD2^4A/4A^* mice (E), disruption of CD2-NHERF-1/2 interaction impairs tip-link maintenance, resulting in mild hair bundle disorganization, shortening of third-row stereocilia, and progressive high-frequency hearing loss.

Of note, residual CD2 staining in *Nherf1^KO/KO^* or *CD2^4A/4A^* stereocilia raised the possibility that CD2 hair bundle localization may be regulated by additional PDZ-containing binding partners such as Whirlin or Harmonin ^27,28^. Our data also imply that stereociliary localization of PCDH15-CD3 and Cadherin-23 may be regulated by NHERF-independent PDZ-PBM interactions. PDZ-PBM interactions have an important role in organizing protein complexes within the hair bundles^27–30^. Several PDZ-domain scaffolding proteins associated with deafness, including Harmonin, Whirlin, and GOPC, interact with different hair cell and stereociliary proteins^28–33^. Thus, our findings further reveal a PDZ-PBM code that controls spatial distribution of specific hair bundle proteins along the length of the stereocilia.

*CD2^4A/4A^* OHCs exhibited normal CD2 localization to the kinocilium and very mild hair bundle defects, indicating the 4A mutation does not significantly affect CD2 function in kinociliary links or hair cell planar polarity during embryonic development^10,13,34^. The more pronounced planar polarity and kinocilium positioning defects in *Nherf1^KO/KO^* compared with *CD2^4A/4A^* suggest that NHERF-1 has additional CD2-independent functions in cochlear development. Since NHERF-1 is highly expressed in supporting cells, the PCP defects may reflect both hair cell-autonomous and non-autonomous functions of NHERF-1 in PCP regulation.

Despite reduced CD2 stereociliary localization, the staircase formation, hair bundle cohesion, and MET were largely normal in *CD2^4A/4A^* OHCs. Thus, the residual amount of CD2 in stereocilia, together with the CD1 and CD3 isoforms, is sufficient to sustain the formation of transient links and tip links and the developmental onset of MET. Remarkably, adult *CD2^4A/4A^* mice exhibited high-frequency hearing loss, accompanied by the degeneration of the third row of stereocilia, whose height is known to be maintained by hair cell MET activity^35,36^. These findings suggest that the 4A mutations impair CD2-dependent tip-link maintenance of mature hair cells. Interestingly, tip link repair in adult hair cells was shown to be initiated by formation of transient PCDH15-PCDH15 links which are then replaced by the mature PCDH15-CDH23 links^36^. Unfortunately, our attempts to immunolocalize CD2 in adult hair cells using existing reagents were unsuccessful. Nevertheless, our data suggest that the 4A mutation, and by inference disrupted CD2-NHERF-1/2 interactions, results in defective tip-link maintenance and repair in adult hair cells. Although NHERF-1 is down-regulated and expressed at low levels in adult hair cells ^37^, NHERF-2 is highly expressed and localized to the hair bundle in mature IHCs and OHCs. Thus, it is plausible that NHERF-2 maintains a reserve pool of CD2 in stereocilia of mature hair cells for rapid tip link repair. NHERF-2 knockout mice exhibited normal hearing and mild degeneration of the third-row stereocilia in OHCs^16^, suggesting genetic compensation by NHERF-1 in adult *Nherf2^KO/KO^* hair cells. In the future, it would be interesting to analyze hair bundle maintenance in adult hair cells deficient for both *Nherf1* and *Nherf2*.

We show that the CD2-NHERF-1/2 interaction is regulated by phosphorylation of the CD2 cytoplasmic tail. Among the several phosphorylation sites within and adjacent to the CD2 PBM, the threonine at the −2 position (T1781) is crucial for binding, predicted to form a hydrogen bond with H59 in the ligand-binding groove of the NHERF-1 PDZ1 domain. As both T1781A and T1781E disrupted this hydrogen bond and greatly reduced CD2-NHERF-1 interaction, we expect that phosphorylation of T1781 to have a similar effect. Indeed, there are other known examples of phosphorylation at the −2 Serine/Threonine of class I PBMs inhibiting PDZ binding^20,38^. Notably, a phosphomimetic mutation of the −2 Threonine in the PBM of the β2-adrenergic receptor strongly reduced its binding to NHERF-1 PDZ1^39^. Thus, although we were not able to demonstrate CD2 T1781 phosphorylation *in vivo*, our results together with previous findings suggest that T1781 phosphorylation may negatively regulate CD2-NHERF-1 binding, thereby reducing CD2 stereociliary localization. In addition, our results suggest that phosphorylation of serine residues distal to the PBM, including S1771 and S1776, further modulates CD2-NHERF-1 binding affinity.

Conversely, the accessibility of NHERF-1 PDZ1 has also been shown to be negatively regulated by phosphorylation in other systems ^40^. Of note, the levels and localization of CD2 in the developing hair bundle correlate with those of NHERF-1. At P0, there are high levels of both proteins in the hair bundle to meet the demands of forming numerous transient lateral links, kinociliary links, and tip links. By P5, both NHERF-1 and CD2 levels in the hair bundle are significantly downregulated, coinciding with the resorption of transient links and kinociliary links ^3^. It is tempting to speculate that these two proteins are coordinately regulated by phosphorylation to achieve the proper level and localization in the developing hair bundle. More broadly, we further propose that phosphorylation is a general mechanism to fine-tune the PDZ-PBM code for specific hair bundle protein interactions. The identity of the CD2 kinase(s) remains to be determined. Modulating the activity of the responsible kinases may be an effective therapeutic strategy for promoting tip-link repair, thereby preventing permanent hearing loss.

## Methods

### Plasmids and expression constructs

Plasmids encoding the transmembrane and cytoplasmic domains (TM+Cyto) of CD1, CD2, or CD3 fused to EGFP were gifts from Dr. Raj Ladher ^13^. The CD2 (TM+Cyto) coding sequences were cloned into pEGFP-N1 and used to generate the T1781A, T1781E, and 4A mutant constructs by site-directed mutagenesis. All mutations were confirmed by Sanger sequencing. The His-MBP-CD2-Cyto plasmid was generated by cloning the fragment encoding the mouse CD2-Cyto into pMAL-c2 ^41^ using the BamHI/NotI restriction sites. GST-NHERF-1 and −2 constructs were generated by cloning full-length cDNAs for NHERF-1 (NCBI: NM_012030.2) and NHERF-2 (NCBI: BC065778.1) into pGEX-6P-1 (Addgene) using the BamHI and EcoRI restriction sites. Fragments containing individual PDZ domains of NHERF-1 were cloned into pGEX-4T-1 using NEBuilder® HiFi DNA Assembly Cloning Kit (E5520S, New England Biolabs).

### Transfections, immunoprecipitation, and mass spectrometry

HEK293T cells were maintained in high-glucose Dulbecco’s modified Eagle’s medium (DMEM) (10313-021, Gibco) supplemented with 10% fetal bovine serum (FBS). 5 × 10^6 cells were transfected with 10 µg of plasmid using the calcium-phosphate transfection method ^42^. At 24–36 h after transfection, cells were lysed in lysis buffer containing 0.2 M NaCl, 5% glycerol, 1% Triton X-100, 2 mM EDTA, 20 mM Tris-HCl, pH 7.5, and Complete® protease and PhosStop phosphatase inhibitors (11697498001 and 04096837001, Roche). Lysates were incubated on ice for 15 min and then centrifuged at 13,000 rpm for 10 min. The supernatants were collected for downstream applications. To identify phosphorylation sites within the C-terminal region of CD2, lysates with overexpressed EGFP-CD2 (TM+Cyto) were immunoprecipitated using anti-GFP agarose beads (ChromoTek, gta) and digested with trypsin (Pierce, 90057). The resulting peptides were desalted using ZipTip-C18 tips (ZTC18S, Millipore) according to the manufacturer’s instructions and submitted for MS/MS analysis as previously described ^43^.

### Generation of and validation of phospho-specific CD2 antibodies

Custom anti-CD2-pS1771 rabbit polyclonal antibodies were produced by Genscript BioTech. Rabbits were immunized with the phosphorylated peptide EPKKpSPGANSEGYC, and phospho-specific antibodies were purified by pre-adsorption with non-phosphorylated peptides followed by affinity purification using phosphorylated peptides. Purified antibodies were validated by Western blot analysis of wild-type and phospho-deficient EGFP-CD2 (TM+Cyto) expressed in HEK293T cells, using bacterially expressed, unphosphorylated MBP-CD2 as an additional negative control.

### Purification of His-MBP fusion proteins and GST fusion proteins

BL-21 (DE3) RIPL bacterial cells were transformed with expression vectors and grown in Terrific Broth medium in the presence of antibiotics. Cultures were induced with 0.5 mM IPTG and grown at 37 °C for 3.5 hours (for His-MBP-CD2cyto), or with 0.25 mM IPTG and grown at 16 °C overnight (for GST fusion proteins). Bacteria were pelleted, resuspended in 30 mL of lysis buffer (20 mM Tris pH 8.0, 500 mM NaCl, 1 mg/ml of hen egg lysozyme, and Complete® protease inhibitors (11697498001, Roche) and lysed by sonication. Lysates were centrifuged at 80,000×*g* for 30 min and subjected to affinity purification.

For the His-MBP fusion protein, the supernatant was loaded onto a Ni-NTA gravity column (88221, Thermo Scientific) and washed with Buffer A (20 mM Tris, 500 mM NaCl, pH 7.5 with 15mM imidazole). Bound proteins were eluted with 300mM imidazole, dialyzed into PBS, and further purified by anion-exchange chromatography. Eluted protein was coupled to NHS-activated Sepharose (17090601, Cytiva) according to the manufacturer’s instructions.

For GST fusion proteins, the supernatant was then loaded onto a gravity column with glutathione agarose (catalog no. 16100, Pierce) and washed with 10 Column Volumes (CV) of Buffer A (20 mM Tris, 500 mM NaCl, pH 8.0) to remove nonspecific binding.

### Affinity purification of CD2-binding proteins and mass spectrometry analysis

Immobilized His-MBP and His-MBP-CD2 proteins were equilibrated with a wash buffer containing 0.1 M NaCl, 4 mM EDTA, 0.01% Triton X-100, and 10 mM Tris-HCl, pH 7.5 supplemented with Complete® protease inhibitors (11697498001, Roche). P0 mouse brain/tissue samples were homogenized in lysis buffer containing 0.1 M NaCl, 10 mM Tris-HCl, pH 7.5, 4 mM EDTA, 1% Triton X-100, 0.3 mM DTT, Complete® protease and PhosStop phosphatase inhibitors (11697498001 and 04096837001, Roche). Lysates were centrifuged at 15000 rpm for 10 minutes at 4°C. Tissue lysates were first pre-adsorbed on a His-MBP column followed by affinity purification using a His-MBP-CD2 column. After washing with 10 column volumes (CV) of wash buffer, bound proteins were eluted with 0.1 M glycine/50 mM HCl from both columns. Proteins were digested with trypsin (90057, Pierce). The resulting peptides were desalted using ZipTip-C18 tips (ZTC18S, Millipore) according to the manufacturer’s instructions and submitted for MS/MS analysis as previously described ^43^.

Peptides detected from samples bound to His-MBP-CD2 but not His-MBP were analyzed and plotted as a function of scaled Mascot score versus emPAI using a custom Python-based visualization workflow. For each fraction, protein abundance, represented by emPAI, was plotted on the x-axis, and the scaled Mascot score (representing -10log_10_(P), where P is the random match probability) was plotted on the y-axis. A Mascot Score of 600 was used as a threshold to highlight the highest-scoring proteins. Highest score proteins, NHERF-1 (Mascot Score = 1032, emPAI =2.2), NHERF-2 (Mascot Score = 634, emPAI =1.41) and PCDH15-CD2 (Mascot Score = 1133, emPAI = 0.24) were mapped on the plot. Proteins of interest such as ezrin (Mascot Score = 120, emPAI = 0.36), radixin (Mascot Score = 118, emPAI = 0.22), and moesin (Mascot Score = 119, emPAI = 0.31) were also highlighted in distinct colors.

### GST pull-down assays and Western blot analysis

For each pull-down reaction, approximately 5-10 μg of Immobilized GST–NHERF-1 and GST–NHERF-2 were incubated overnight at 4 °C with extracts from HEK293T cells overexpressing pEGFP-CD1/2/3 (TM+Cyto) or mutant CD2 constructs (T1781A, T1781E, or 4A). Following four washes with wash buffer (0.2 M NaCl, 0.01% Triton X-100, 2 mM EDTA, 20 mM Tris, pH 7.5, and Complete® protease and PhosStop phosphatase inhibitors (11697498001 and 04096837001, Roche)). Bound proteins were eluted with 15 mM glutathione and separated by SDS-PAGE using Bio-Rad’s Mini Protean, transferred to a nitrocellulose membrane, and analyzed by Western blotting, Coomassie and/or Ponceau staining. Primary and secondary antibodies used for immunoblotting are listed in Table 1. Infrared fluorescence on membranes was visualized using the Li-COR Odyssey® CLx Imaging System. Coomassie staining was performed using QC Colloidal Coomassie Stain. Western blot band intensities were quantified using Fiji/ImageJ (National Institute of Health). Bound protein signals were normalized to those of the corresponding extracts, with bound levels of wild-type EGFP-CD2(TM+Cyto) set to 100%.

**Table 1.** List of Antibodies.

| Antibody | Concentration | Vendor | Catalog# |
| --- | --- | --- | --- |
| Alexa-conjugated phalloidin | 1:200 | Thermo Fisher | A12379, A12380, A12381, A22287 |
| Alexa-conjugated secondary antibodies | 1:500 | Thermo Fisher | A11036, A11029, A11057, A11077 |
| IRDye 680 LT Goat anti-Mouse IgG | 1:20000 | LICORbio | 926-68020 |
| IRDye 800CW Goat anti-Rabbit IgG | 1:20000 | LICORbio | 926-32211 |
| Anti-GFP | 1:2000 WB | Invitrogen | A11122 |
| Anti-acetylated tubulin | 1:400 | Sigma | T6793 |
| Anti-EBP50/NHERF-1 | 1:100 | Proteintech | 29771-1-AP |
| Anti-Eps8 | 1:200 | BD | 610143 |
| Anti-PCDH15-CD2 | 1:200 | Dr. Uli Mueller, JHU | Reference 10 |
| Anti-Cadherin-23 | 1:300 | Dr. Uli Mueller, JHU | Reference 19 |
| Anti-PCDH15 (PB811) | 1:200 | Dr. Bechara Kachar, NIH | Reference 18 |
| Anti-PCDH15-CD2 (PB860/61/62) | 1:200 | Dr. Bechara Kachar, NIH | Reference 18 |
| Anti-PCDH15-CD2-pS1771 | 10ug/ml IF | Custom-made by Genscript | This study |

### Mice

*Nherf1^KO^* mice (Strain #:012862) were ordered from The Jackson Laboratory ^17^. The *CD2^4A^* knock-in allele was generated by the Genetically Engineered Murine Model Core Facility at the University of Virginia using CRISPR-assisted genome editing in mouse zygotes, as previously described ^44^. The guide RNA (crRNA), tracrRNA, ssODN, and Cas9 protein were purchased from IDT (Coralville, Iowa). Founders were backcrossed with wild-type, *Cdh23^Ahl+^* B6SJL mice 3 times, and offspring were used for intercrosses. The following genotyping primers were used: *Nherf1^KO^*, 5’-ttcggcctcattctggtc-3’ and 5’-cgccttcttgacgagttctt-3’; *Nherf1^+^*, 5’-ttcggcctcattctggtc-3’ and 5’-gagaagggtccaaatggcta-3’. *CD2^4A^*, 5’-agcccctggagccaatgctgc-3’ and 5’-ttgctatagacaagtgtcactg-3’; *CD2^+^*, 5’-atctccaggcgcaaactccgaa-3’ and 5’-ttgctatagacaagtgtcactg-3’. Animal care and use were conducted in accordance with National Institutes of Health guidelines and approved by the Animal Care and Use Committee at the University of Virginia.

### Immunohistochemistry and Fluorescence Microscopy

Skulls from P0, P5, and P6 mice were dissected and fixed in 4% paraformaldehyde for 1 hour at room temperature, then washed three times in PBS. For whole-mount immunostaining, cochleae were dissected from the temporal bones, and the right cochlea from a control mouse was paired with the left cochlea from a mutant mouse, or vice versa. Each control–mutant pair was processed together under identical conditions. Samples were blocked for 1 hour at RT in PBS containing 0.1% Triton X-100, 5% heat-inactivated sheep serum, and 0.05% sodium azide, then incubated with primary antibodies for 18–48 hours at 4 °C. After three washes in PBS containing 0.1% Triton X-100 (PBS-T), samples were incubated with secondary antibodies and phalloidin for 2 h at RT. Samples were then washed in PBS-T, post-fixed in 4% paraformaldehyde for 15 min at RT, and washed twice more in PBS-T before being flat-mounted in Fluoromount-G.

Immunofluorescence images were acquired using a DeltaVision deconvolution microscope or a Leica SP8 confocal microscope. CD2 localization to the kinocilium was imaged using a Leica STELLARIS 8 tauSTED super-resolution microscope at the Advanced Microscopy Facility, and phospho-S1771-CD2 staining images were acquired using a Nikon SoRa spinning disk confocal microscope. Control and mutant samples from the same experiment were imaged using identical acquisition settings. Multiple images were acquired from the basal, middle, and apical regions of the cochlear duct, corresponding to 10%, 50%, and 80% of the total cochlear length, respectively, to assess any tonotopic changes in staining intensity and hair bundle phenotypes. Images were processed using Fiji/ImageJ (National Institutes of Health) and Photoshop (Adobe) to adjust brightness, contrast and gamma.

For quantification of staining intensity, identical brightness and level adjustments were applied to representative control and mutant images. Regions of interest (ROIs) were manually drawn around individual hair bundles using the freehand selection tool, and the mean fluorescence intensity of the protein of interest was normalized to that of phalloidin staining in the same ROI. To assess tonotopic changes in staining intensity, normalized staining intensity in individual hair cells was plotted relative to the mean ratio of the control basal region, which was set at 100%. Statistical significance was determined using a mixed-effects model or with multiple comparisons testing to assess differences between cochlear regions and genotypes, with p < 0.02 considered statistically significant.

### SEM

SEM sample preparation was performed based on a previously published protocol ^45^. Briefly, temporal bones were removed from the skull of P30 mice, perfusion-fixed immediately, and then fixed in 4% paraformaldehyde and 2.5% glutaraldehyde overnight at 4°C. P30 samples were decalcified in 10% EDTA for 3-4 days. Cochleae were carefully dissected in dissection buffer (0.9% NaCl, 0.05 mM HEPES pH 7.2, 2 mM CaCl₂, 0.5 mM MgCl₂) with systematic removal of surrounding bone, stria vascularis, Reissner’s membrane, and tectorial membrane to fully expose the organ of Corti. Dissected samples were transferred to secondary fixative (2.5% glutaraldehyde, 4% paraformaldehyde in dissection buffer) and incubated at 4°C overnight. Following secondary fixation, samples were washed in dissection buffer (3 × 5 min) and dehydrated through a graded ethanol series on ice (15-95% in 15% increments, then 2 × 100% ethanol, 10 minutes each) under a fume hood. Samples were further processed using hexamethyldisilazane (HMDS; 440191, Sigma-Aldrich) treatment with increasing concentrations (30-90% in 30% increments, then 3 × 100% HMDS, 15 minutes each), followed by overnight drying under a fume hood. Dried samples were mounted on specimen stubs, sputter-coated with gold-palladium to 8 nm thickness, and imaged using a Jeol-JSM-IT-800HL FE-SEM.

### Whole-cell voltage-clamp electrophysiology recordings

We analyzed mechanotransduction (MET) currents in OHCs from the mid-apical turn of *CD2^4A/+^* and *CD2^4A/4A^* mice at P5-P7 as previously described ^46,47^. Hair bundles were mechanistically stimulated using either stepwise stereociliary deflections with a stiff glass probe at P5-P6 or sinusoidal fluid-jet stimulation of apical OHC hair bundles at P6-P7. During recordings, apical perfusion was used to apply artificial perilymph (in mM): 144 NaCl, 0.7 NaH2PO4, 5.8 KCl, 1.3 CaCl2, 0.9 MgCl2, 5.6 glucose, and 10 H-HEPES, pH 7.4. Borosilicate glass pipettes with filaments were pulled with a P-2000 pipette puller (Sutter, CA), and the tips were polished with an MF-930 microforge (Narishige, Japan) to resistances of 2–3 MΩ. Mechanical stimulation of hair bundles was performed with a stiff glass probe mounted on a piezoelectric stack actuator (Thorlabs, Newton, NJ) or a fluid Jet from a glass pipette, which is produced by a 27-mm-diameter piezoelectric disc. For stiff glass probe stimuli, the actuator was driven by voltage steps (10 ms) that were low-pass filtered at10 kHz with a 900CT eight-pole Bessel filter (Frequency Devices) to diminish the resonance of the piezo stack. The tip of the probe was fire-polished to fit the shape of OHC bundles. For Fluid Jet stimuli, a 40 Hz sinusoidal voltage was applied and filtered at 1.0 kHz. Whole-cell recordings were carried out with an EPC 10 USB patch-clamp amplifier (HEKA, Germany), and signals were sampled at 100 kHz with a Bessel 10 kHz filter. The signals were offline filtered at 5kHz. Hair cells were voltage-clamped at −84mV with intracellular solution (in mM): 140 mM CsCl, 1 mM MgCl2, 1 mM BAPTA, 2 mM Mg-ATP, 0.3 mM Na-GTP, and 10 mM H-HEPES, pH 7.2. Uncompensated series resistance was less than 5 MΩ.

Data analysis was performed using Excel (Microsoft) and Igor Pro 7 (WaveMetrics, Lake Oswego, OR). Transduction current/displacement curves (I/X) and open probability/ displacement curves (Po/X) were fitted with the following double Boltzmann equation:

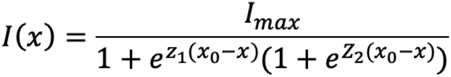

Where Z1 and Z2 are the slope factors, and X_0_ represents the set point.

### Auditory Brainstem Response (ABR) Measurements

ABR tests were performed in a sound-attenuating chamber (Med-Associates, product no. ENV-022MD-WF) using a Tucker-Davis Technologies (Alachua, FL) system as previously described ^48^. Briefly, mice were anesthetized with a single intraperitoneal injection of ketamine hydrochloride, 87.5 mg/kg (Covetrus, NDC 11695-0703-1), and xylazine hydrochloride, 12.5 mg/kg (Dechra, NDC 17033-099-05). Thresholds were determined from responses to 1024 stimulus presentations at 8, 11, 16, 22, and 32 kHz. Stimulus intensity was adjusted in 5 dB increments until a reproducible waveform was identified. If no waveform was detected at the maximum transducer output, the threshold was assigned as 5 dB above the maximum output. Longitudinal ABR tests were performed in *CD2^4A/4A^* and *CD2^WT^* mice at 1, 2.5, and 5 months of age. ABR analysis of *CD2^4A/4A^* mice included both males and females. Data are presented as sound pressure levels in dB SPL. Statistical significance was assessed using two-way ANOVA with multiple comparisons testing, and graphs were generated using GraphPad Prism.

### Hair bundle orientation and kinocilium index quantifications

Hair bundle orientation and kinocilium positioning were quantified as previously described^49^. Hair bundle orientation was determined by measuring the angle of rotation relative to the horizontal axis of the cochlear duct. Hair bundles oriented toward the lateral edge of the cochlear duct were defined as 0°, whereas HBs oriented parallel to the rows of hair cells, in either direction, were defined as 90°. Kinocilium positioning within the hair bundle was analyzed by calculating the kinocilium index, as previously described ^49^. Briefly, distances from the base of the kinocilium to each end of the HB were measured in Fiji and designated as x and y. The kinocilium index was calculated as the ratio of the longer distance to the shorter distance (x/y). A kinocilium index greater than 1.5 was considered indicative of an off-center kinocilium. Statistical significance was determined using a mixed-effects model ^50^ to assess differences between cochlear regions and genotypes, with p < 0.02 considered statistically significant.

## Acknowledgments

We thank Dr. Bechara Kachar (National Institute of Deafness and Communication Disorders) for sharing reagents and critical reading of the manuscript, Drs. Jung-Bum Shin, Elias Spiliotis, Karen Hirschi and Lukas Tamm for sharing reagents and equipment, Dr. Daniel Grigsby, Dr. Sihan Li and Wenxia Li for technical assistance. This work used a Leica Stellaris 8 tauSTED Super-Resolution Microscope and a Jeol-JSM-IT-800HL FE-SEM in the Advanced Microscopy Facility (University of Virginia, RRID: SCR_018736). This research was supported by a Fellowship grant from the American Otological Society (to A. Adylkhan) and National Institutes of Health grants R01DC013773 and R01HD107872 (to X. Lu), R01DC023325 (to U. Mueller), the Pinn Scholar Award from the University of Virginia School of Medicine (to X. Lu), and grants from Nazarbayev University (to A. Andreeva).

---

**Figure S1.**
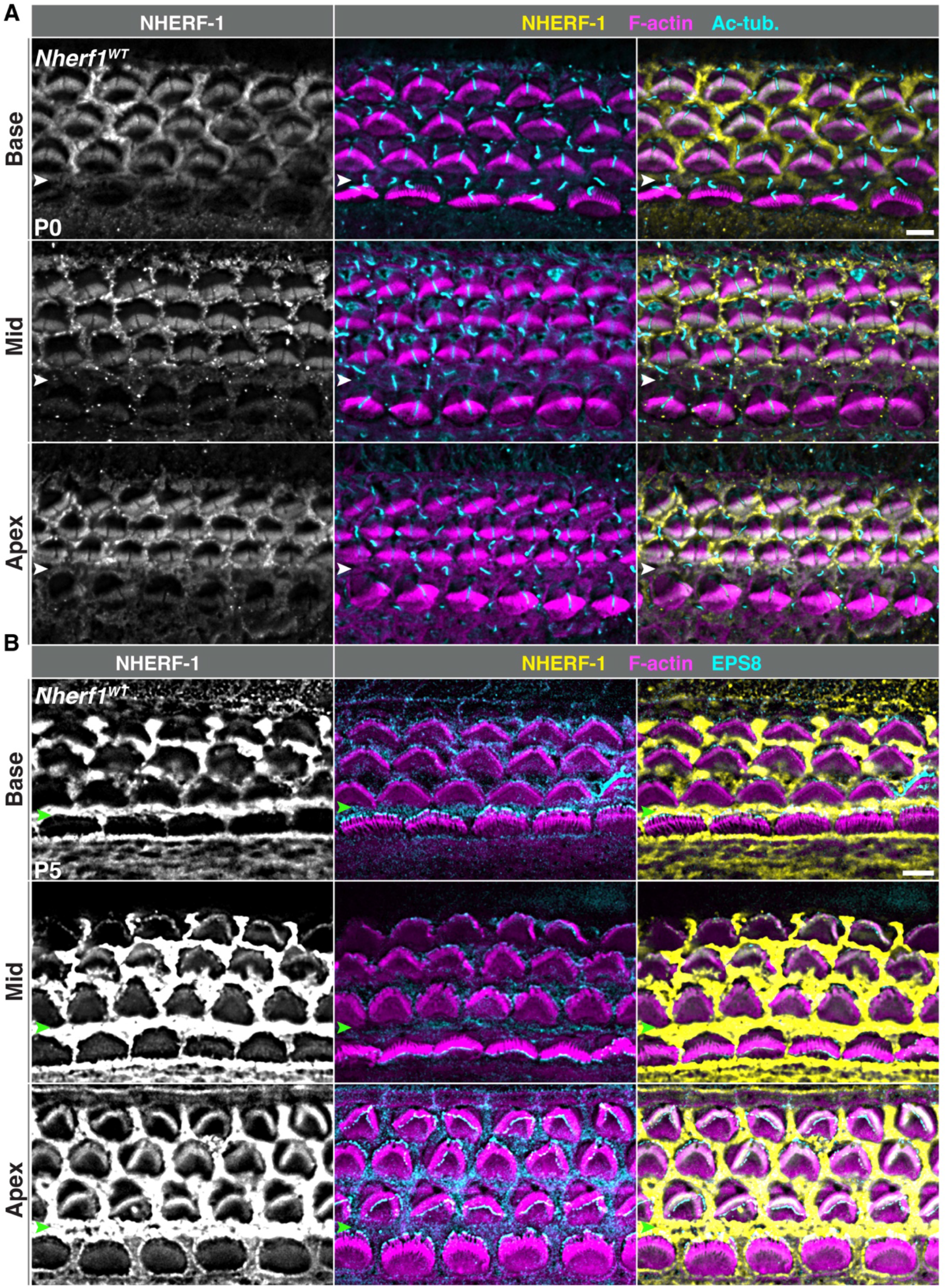
NHERF-1 localization in the organ of Corti at P0 and P5. **(A)** Representative images of whole-mount organ of Corti surface preparations from the basal, middle, and apical turns stained for NHERF-1 (yellow), F-actin (magenta), and acetylated tubulin (Ac-tub, cyan) at P0. **(B)** Representative images of whole-mount organ of Corti surface preparations from the basal, middle, and apical turns stained for NHERF-1 (yellow), F-actin (magenta), and EPS8 (cyan) at P5. Arrowheads mark the IPC row. Scale bars, 6 μm.

**Figure S2.**
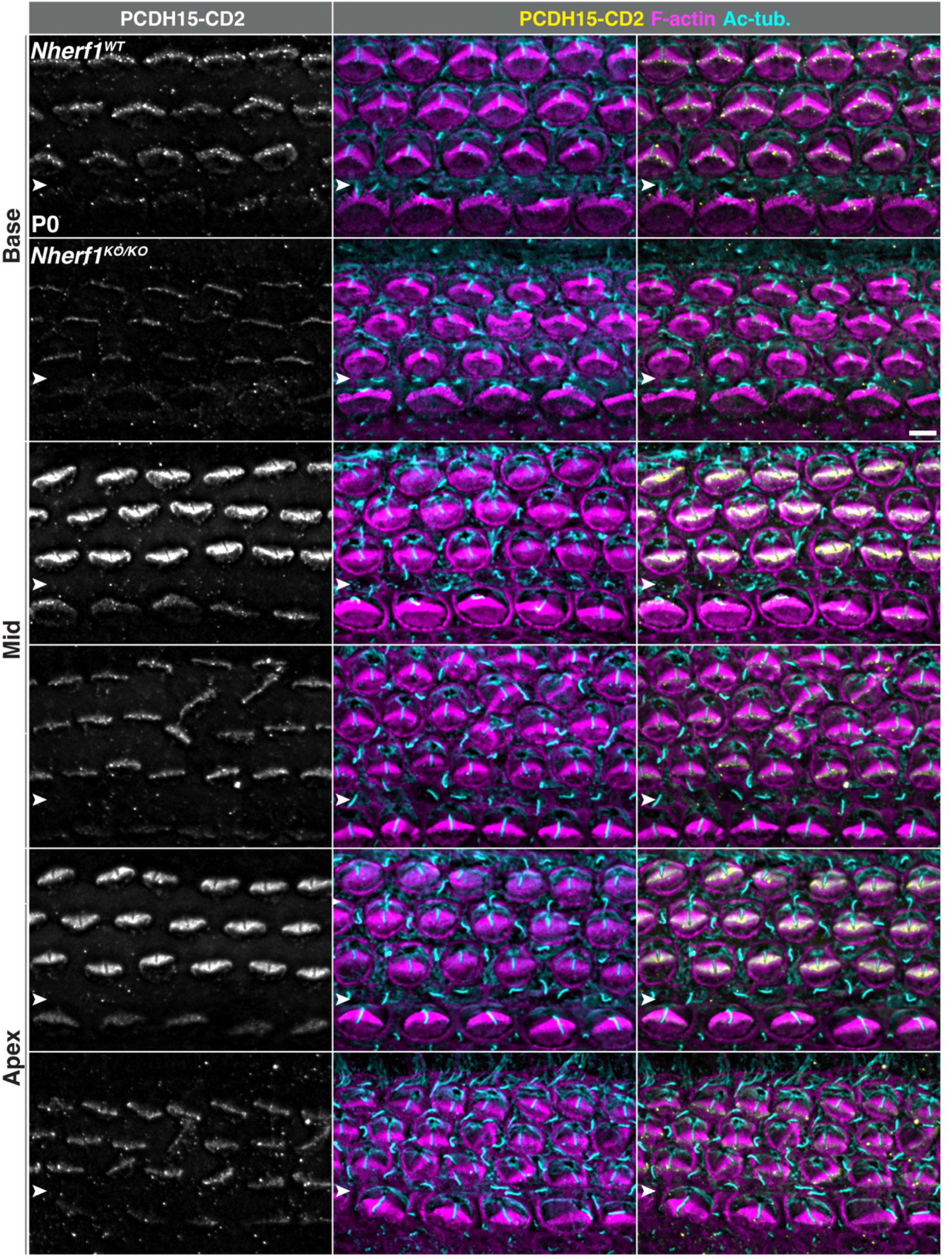
Reduced stereociliary localization of PCDH15-CD2 in *Nherf1^KO/KO^* OHCs at P0. Representative images of whole-mount organ of Corti surface preparations from the basal, middle, and apical turns stained for PCDH15-CD2 (yellow), F-actin (magenta), and acetylated tubulin (Ac-tub, cyan). White arrowheads mark the IPC row. Scale bar, 6 μm.

**Figure S3.**
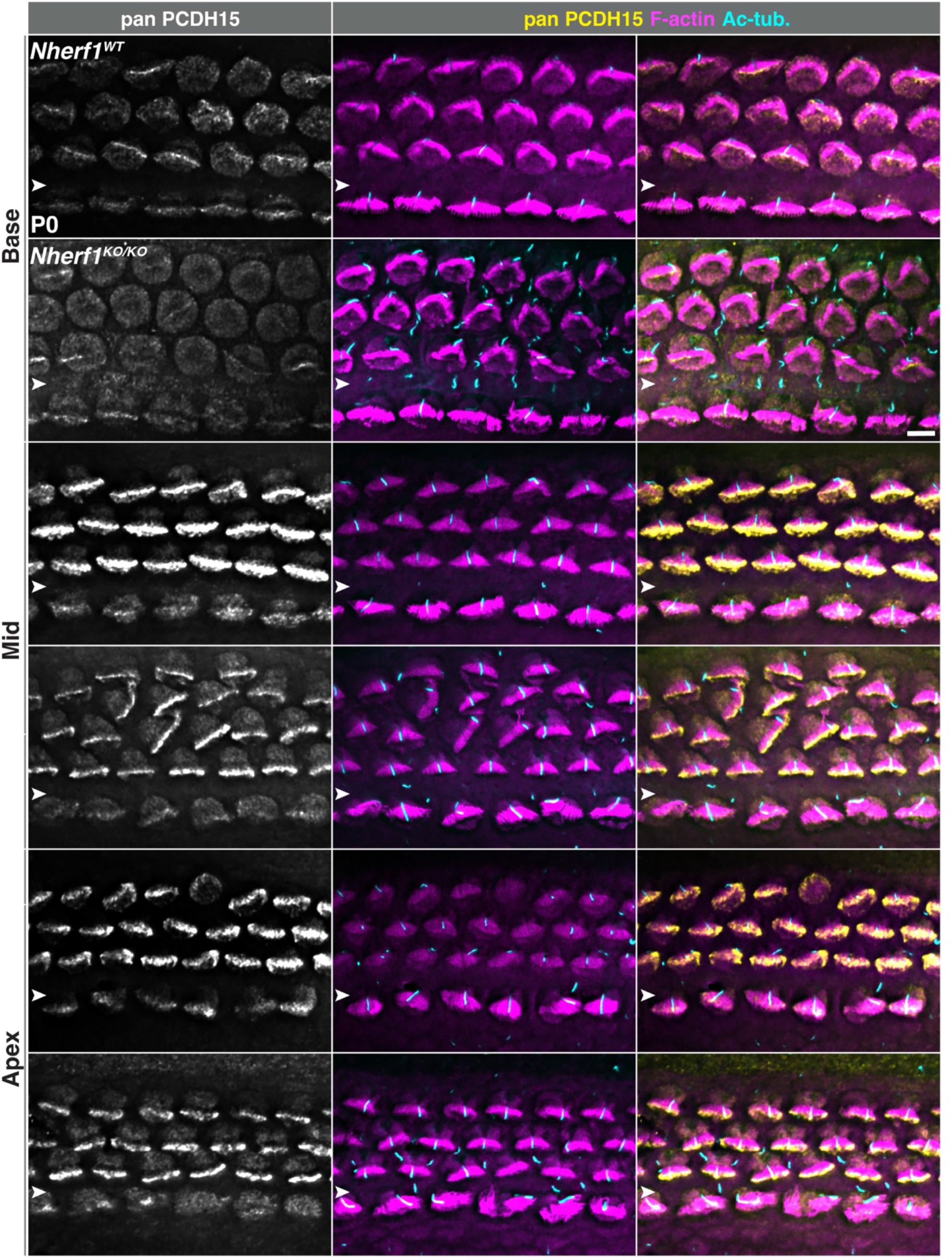
Stereociliary localization of PCDH15 in *Nherf1^KO/KO^* mice at P0. Representative images of whole-mount organ of Corti surface preparations from the basal, middle, and apical turns stained for all PCDH15 isoforms (yellow), F-actin (magenta), and acetylated tubulin (Ac-tub, cyan). White arrowheads mark the IPC row. Scale bar, 6 μm.

**Figure S4.**
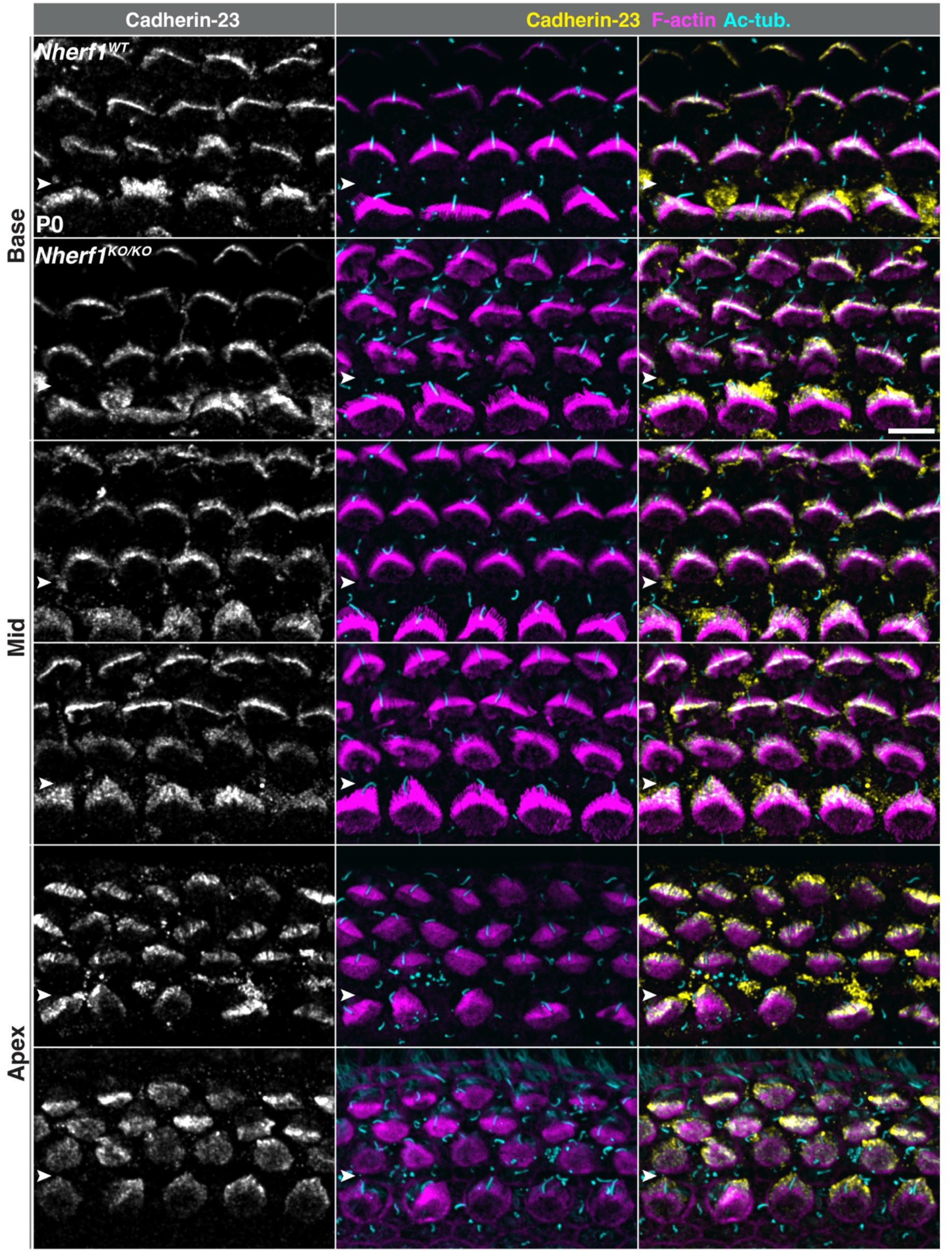
Stereociliary localization of Cadherin-23 is unchanged in *Nherf1^KO/KO^* mice at P0. **(A)** Representative images of whole-mount organ of Corti surface preparations from the basal, middle, and apical turns stained for Cadherin-23 (yellow), F-actin (magenta), and acetylated tubulin (Ac-tub, cyan). White arrowheads mark the IPC row. Scale bar, 6 μm.

**Figure S5.**
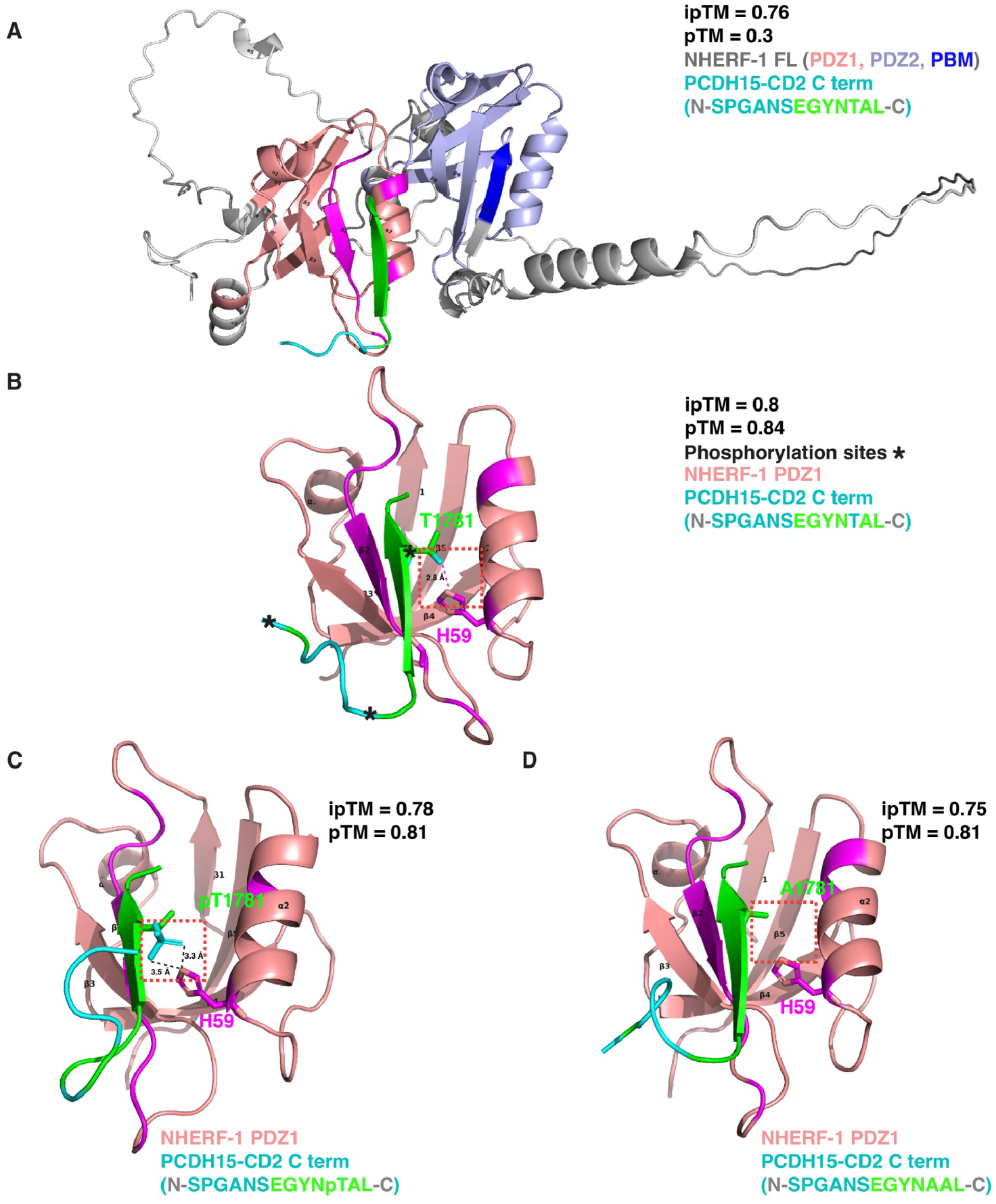
AlphaFold structural modeling of CD2-NHERF-1 interaction. **(A)** Predicted complex between full-length mouse NHERF-1 and the CD2 cytoplasmic tail (cyan and green). The CD2-PBM (green) nestles into the binding groove of the PDZ1 domain (pink), and the NHERF-1 PBM (dark blue) forms intramolecular interactions with the PDZ2 domain (light blue). **(B)** Predicted complex between isolated NHERF-1 PDZ1 and CD2 cytoplasmic tail. Asterisks mark phosphorylation sites identified in the CD2 cytoplasmic tail. The dashed box indicates the predicted hydrogen bond between H59 in the α2 helix of NHERF-1 PDZ1 and T1781 of CD2-PBM. **(C)** Phosphorylated T1781 of PCDH15-CD2 is predicted to sterically disrupt the T–H hydrogen bond that dictates Class I PDZ binding specificity. **(D)** CD2-T1781A is incapable of forming a hydrogen bond with H59. Modeling was performed using AlphaFold 3-Multimer.

**Figure S6.**
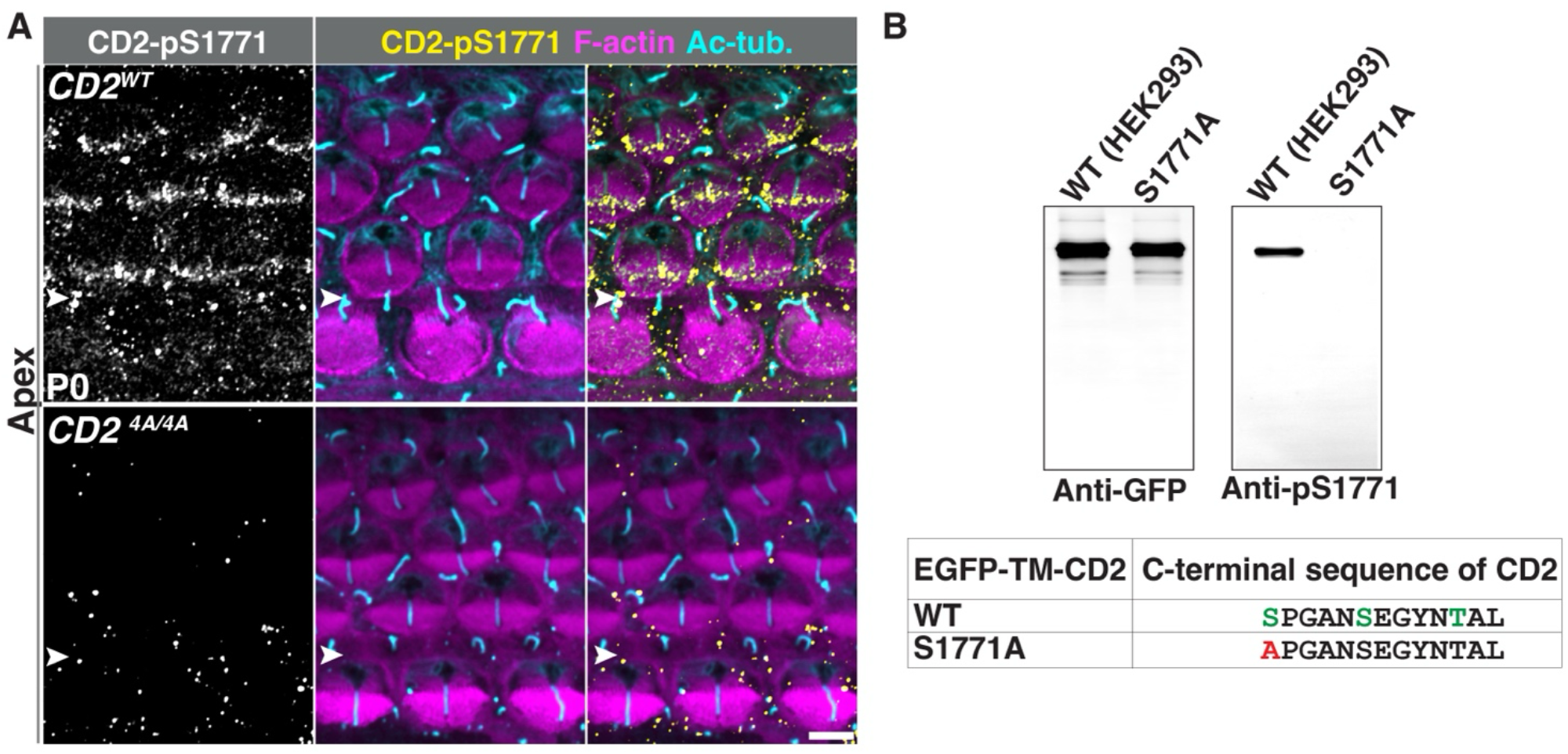
Detection of phosphorylated CD2 in OHC hair bundles *in vivo* at P0. **(A)** Whole-mount surface preparations of the organ of Corti from the apical turn stained for CD2-pS1771 (yellow), F-actin (magenta), and acetylated tubulin (Ac-tub, cyan). White arrowheads mark the IPC row. Scale bar, 2 μm. **(B)** Validation of anti-CD2-pS1771 antibody *in vitro.* Whole-cell lysates from HEK293 cells transfected with wild-type (WT) and the S1771A mutant of EGFP-CD2 (TM+Cyto) were blotted with anti-EGFP (left panel) and anti-pS1771 (right panel) antibodies. Anti-pS1771 antibodies detected WT CD2 but not the S1771A phospho-deficient mutant.

**Figure S7.**
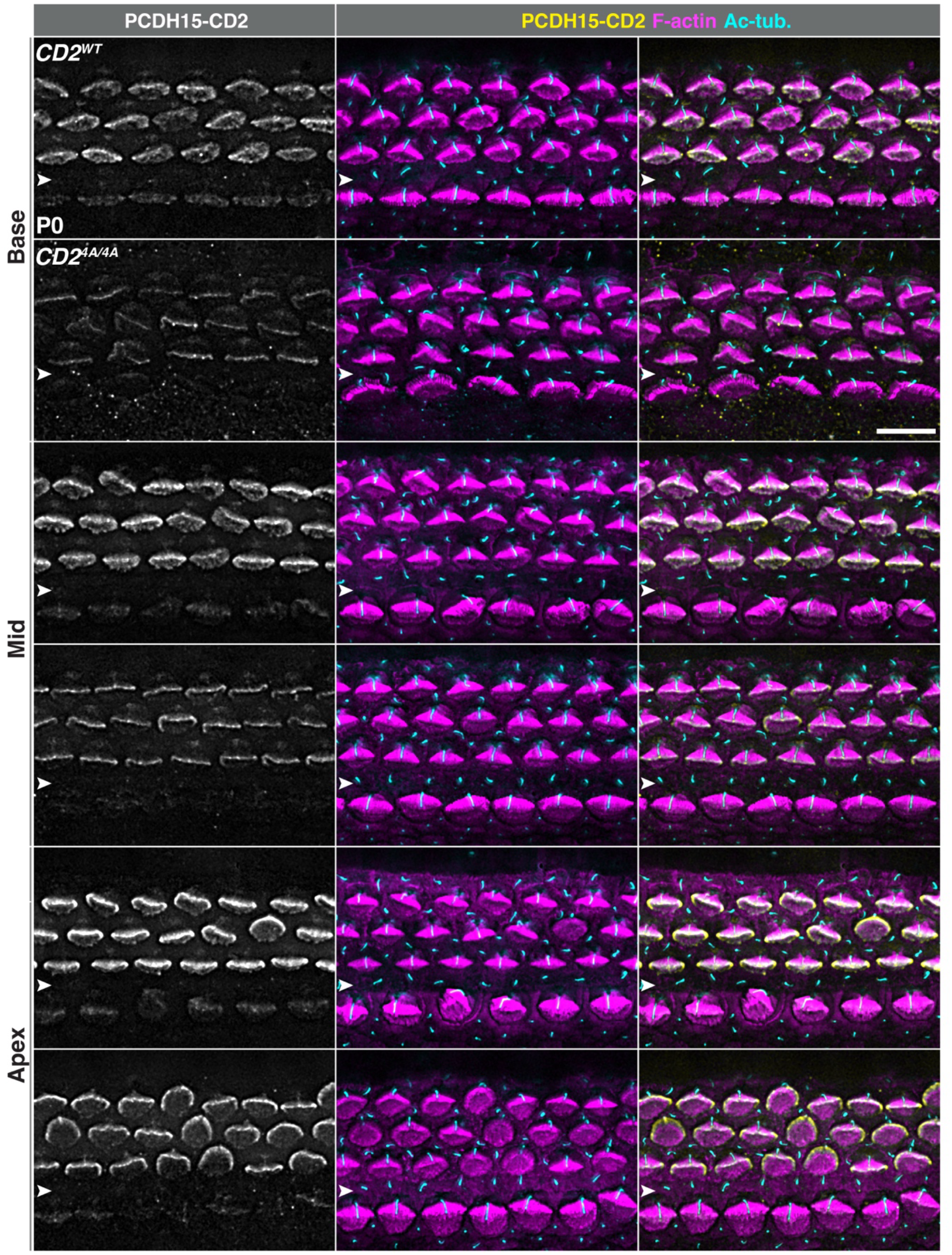
Reduced stereociliary localization of PCDH15-CD2 in the organ of Corti of *CD2^4A/4A^* mice at P0. Representative images of whole-mount organ of Corti surface preparations from the basal, middle, and apical turns stained for PCDH15-CD2 (yellow), F-actin (magenta), and acetylated tubulin (Ac-tub, cyan). White arrowheads mark the IPC row. Scale bar, 6 μm.

**Figure S8.**
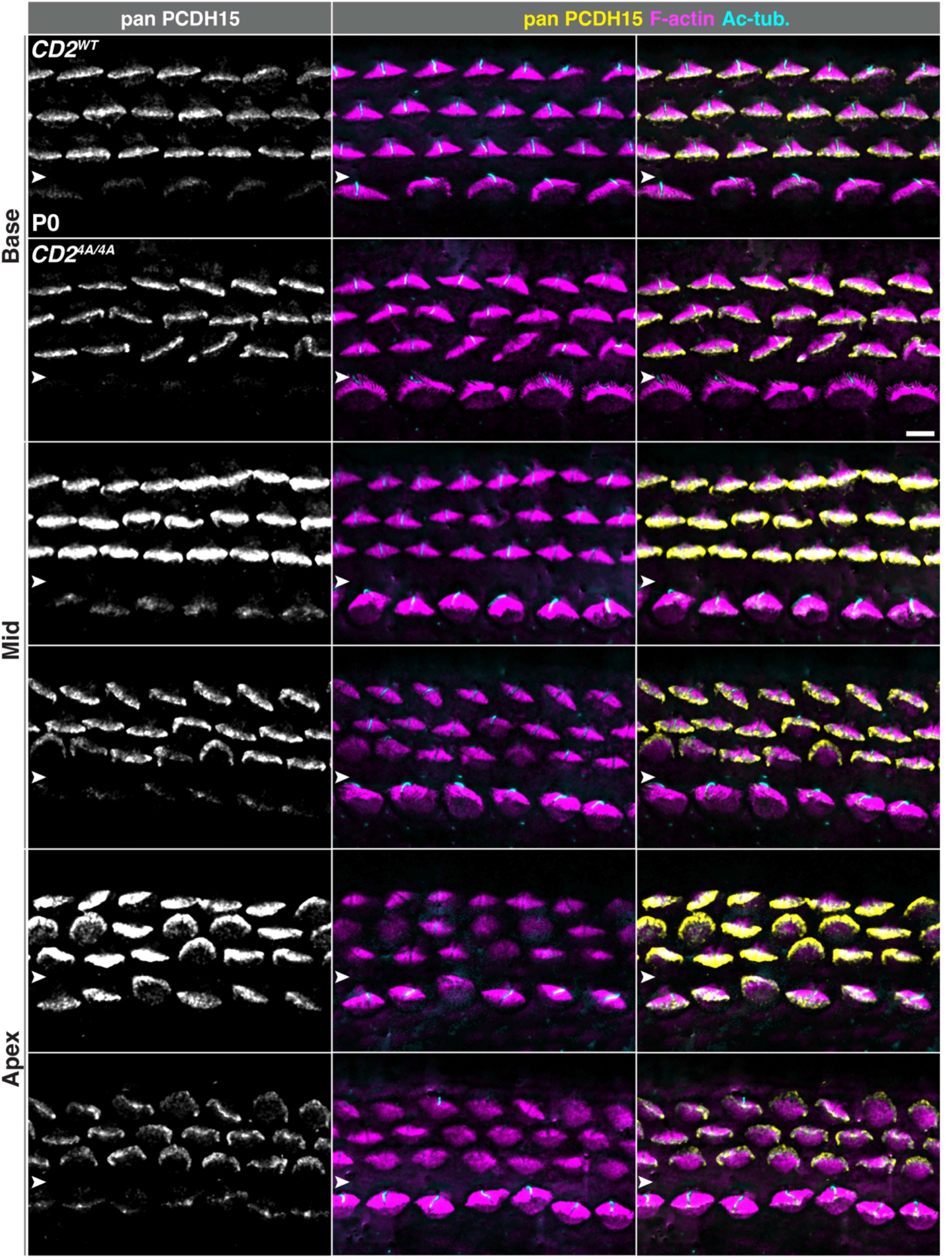
Stereociliary localization of PCDH15 in *CD2^4A/4A^* mice at P0. Representative images of whole-mount organ of Corti surface preparations from the basal, middle, and apical turns stained for all PCDH15 isoforms (yellow), F-actin (magenta), and acetylated tubulin (Ac-tub, cyan). White arrowheads mark the IPC row. Scale bar, 6 μm.

**Figure S9.**
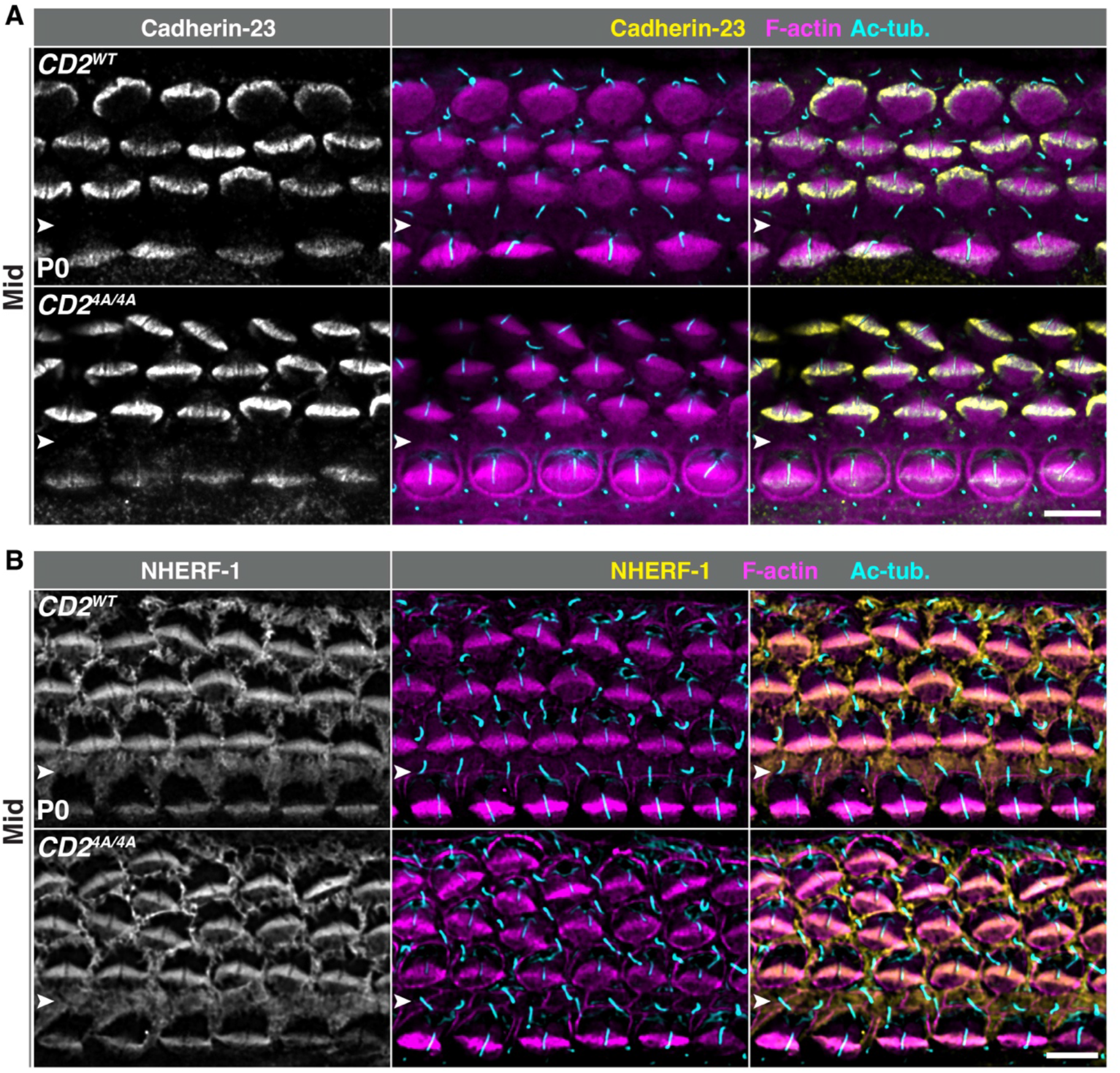
Cadherin-23 and NHERF-1 localization are unchanged in the *CD2^4A/4A^* organ of Corti at P0. **(A, B)** Whole-mount organ of Corti surface preparations from the middle turn stained for Cadherin-23 (A, yellow) or NHERF-1 (B, yellow), F-actin (magenta), and acetylated tubulin (Ac-tub, cyan) at P0. White arrowheads mark the IPC row. Scale bar, 6 μm.

